# Conserved neural dynamics and computations across species in olfaction

**DOI:** 10.1101/2023.04.24.538157

**Authors:** Doris Ling, Elizabeth H Moss, Cameron L Smith, Feiyang Deng, Ryan Kroeger, Jacob Reimer, Baranidharan Raman, Benjamin R Arenkiel

**Author notes:** Department of Anesthesiology and Perioperative Medicine, Oregon Health & Science University, Portland, OR. These authors contributed equally (Doris Ling, Elizabeth H Moss).

## Abstract

Interpreting chemical information and translating it into ethologically relevant output is a common challenge of olfactory systems across species. Are computations performed by olfactory circuits conserved across species to overcome these common challenges? To investigate this, we compared odor responses in the locust antennal lobe (AL) and mouse olfactory bulb (OB). We found that odors activated nearly mutually exclusive neural ensembles during stimulus presentation (‘ON response’) and after stimulus termination (‘OFF response’). Strikingly, ON and OFF responses evoked by a single odor were anticorrelated with each other. ‘Inverted’ OFF responses led to a history-dependent suppression of common ensemble elements, which enhanced contrast between odors experienced close together in time. Notably, odor-specific OFF responses persisted long after odor termination in both AL and OB networks. Taken together, our results reveal key neurodynamic features underlying olfactory computations that are conserved across insect and mammalian olfactory systems.

## INTRODUCTION

Olfactory systems across species have the common requirement of translating chemical information into neuronal activity that then drives behaviors. To achieve this, species ranging from aquatic invertebrates to terrestrial mammals employ neural circuits with remarkable^1,2^. This is striking, given that olfactory sensory organs of vertebrate and invertebrate species sample chemical cues much differently. For example, insect olfaction relies on odor molecules making direct contact with external antennae, whereas mammalian olfaction requires respiration to internalize odor molecules and bring them into contact with the olfactory epithelium^2^. While both vertebrate and invertebrate systems employ large families of seven-transmembrane proteins for sensing odors, in vertebrates they function as G-protein coupled receptors^3^ whereas in insects they act as ligand-gated ion channels^4^. The resulting differences in the structure of sensory input cause markedly different temporal dynamics in odor inputs and odor-evoked neuronal activity^5–7^. In addition, early olfactory circuits of the invertebrate antennal lobe (AL) have been studied predominantly as feed-forward neural networks^8–10^. In comparison, their vertebrate counterparts in the olfactory bulb (OB) receive massive centrifugal feedback projections^11–14^, suggesting that complex processing begins at the first circuit node that receives chemosensory input.

Despite these differences, olfactory systems have common needs that are conserved across species. These include sensitivity and specificity across a complex chemical space, a broad dynamic range, the ability to filter and emphasize different inputs, and the capacity to distinguish distinct odors from complex backgrounds^15–22^. Are these common needs met by shared circuit mechanisms and computational principles across species? Or do different odor sampling methods and olfactory circuit architectures change how odors are represented and processed by vertebrate and invertebrate systems? By comparing neural dynamics in early olfactory processing centers across diverse species, we have identified shared computational principles for how odor information is represented and processed over time – particularly after an odor stimulus ends.

In many sensory circuits, not just olfaction, stimulus-specific information is encoded across space (as neuronal ensembles), and over time (in neural dynamics). Neural responses evoked by even a brief encounter with a stimulus often continue well after stimulus termination as sensory ‘aftereffects’^23–26^. Whether an odor leaves behind a stimulus-specific fingerprint (‘afterimage’), and whether such activity serves a computational purpose that underlies the adaptive processing capabilities of olfactory circuits remains unknown. To address these questions, we compared population-level odor response dynamics across insects and mammals, with the goal of identifying conserved computational principles that guide olfaction. Using *in vivo* electrophysiology in the locust AL, and mesoscale two-photon calcium imaging of the mouse OB, we captured neuronal ensemble responses to passive odor presentations. The complementary approaches for recording odor responses had unique advantages in capturing odor response dynamics with different levels of spatial and temporal resolution in each species. Even despite the different resolutions of these methods and the different cellular sources of odor response signals (pools of dendrites in mice vs. single neurons in locust), we found notable similarities in the patterns of odor response dynamics between mice and locusts. Together, these similarities reveal generalizable functions of early olfactory circuits.

In both mice and locusts, two species with markedly different olfactory system organization, we found that ensembles of neurons activated during odor presentations (ON responses) are different from those that become responsive after stimulus termination (OFF responses). This results in distinct categories of nearly mutually exclusive ON and OFF neuronal response patterns across odors. Strikingly, neuronal ensembles in both locust AL and mouse OB show responses to the onset and the offset of individual odors that are inverted relative to each other (i.e., ON and OFF population response vectors are anticorrelated). We propose that such similar neural dynamics across mouse and locust provide a neural underpinning of two emergent computations shared by both species: contrast enhancement and short-term olfactory memory. Such computations are critical for animals navigating the world, following odor trails, encountering new odors within complex backgrounds, and sensing odors in temporally complex plumes. Together, our results suggest that these fundamental building blocks of olfactory computations are conserved across insect and mammalian olfactory systems, revealing common mechanisms of olfactory processing.

## RESULTS

### OFF responses form anticorrelated odor afterimages across species

To assay population-level odor-evoked activity in the locust antennal lobe (AL), we recorded extracellular electrophysiological activity of AL projection neurons (PN) during passive odor presentations (**Fig. 1A-B**). Notably, the majority of responsive PNs during odor presentations (ON responders) became silent after stimulus termination (**Fig. 1C**). Conversely, neurons that were suppressed during odor presentation became activated after stimulus termination (OFF responders). The mean spike counts across all ON and OFF responsive PNs (**Fig. 1C**, traces) revealed the mutually exclusive nature of responses in these two subsets of projection neurons.

**Fig. 1:**
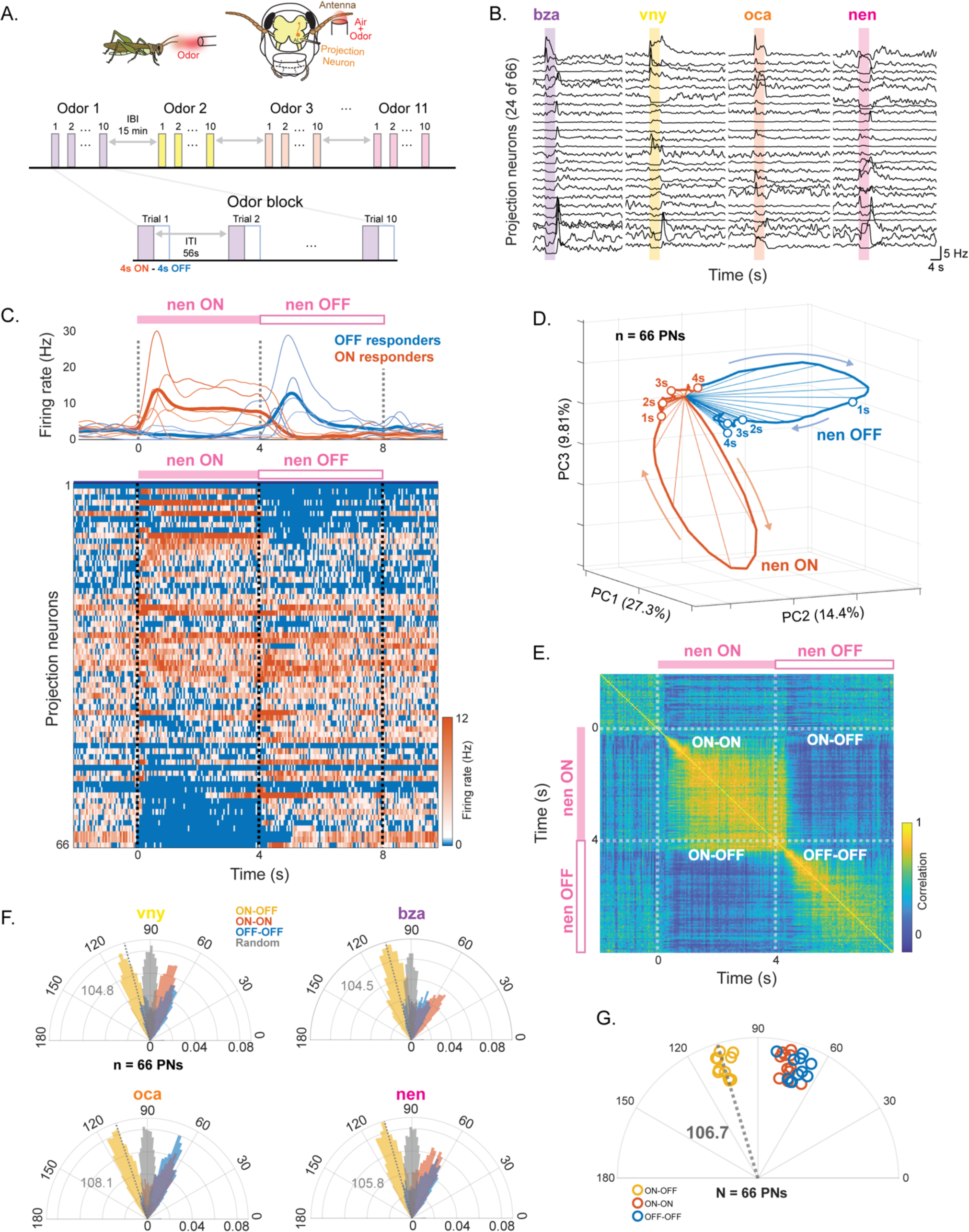
Odor ON and OFF responses in the locust antenna lobe are anticorrelated. **A.** Schematic of the experimental design. (top) Multi-unit extracellular electrophysiological signals were recorded from the projection neurons (PN) in the locust antennal lobe (AL). (bottom) Odors were presented to locusts in blocks of ten trials. A 15-minute gap separated blocks of different odors. **B.** Representative trial-averaged PSTH traces of 24 PNs in response to benzaldehyde (bza), 4-vinyl anisole (vny), octanol (oct), and z-3-nonen-1-ol (nen). **C.** (top) PSTH traces from three ON-responding (orange, thin lines) and OFF-responding PNs (thin blue lines) with respective means (thick lines). (bottom) Trial-averaged responses of 66 PNs to z-3-nonen-1-ol (nen) sorted by ratio of ON to OFF response. **D.** Representative PCA trajectories (thick lines) of trial-averaged PN activity during the 4-sec ON period of nen exposure (orange) and 4-sec OFF period (blue). Spokes (thin lines) represent 200 ms time intervals and open circles mark one second intervals. **E.** Representative temporal correlation of trial-averaged PN activity. **F.** Distribution of angular separation between PN population vectors for four representative odors (vny, bza, oct, and nen). Angular separations are calculated between any ON response vector with other ON response vectors (orange), any OFF response vector with any other OFF response vector (blue), or any ON response vector with any OFF response vector (yellow). Distribution of angular separation between random vectors (grey) are shown as a control. See Figure S1 for other odors. Median values of ON vs OFF angular separation distributions indicated with dashed lines and grey text. **G.** Summary of the median angular separations for all 11 odors showing ON vs ON period median angles (orange), OFF vs OFF period median angles (blue) and ON vs OFF period median angles (yellow). Points corresponding to different odors are offset along the radius for visualization. Mean ON vs OFF angular distance across odors indicated by the dashed grey line.

To qualitatively visualize distinct odor ON and OFF dynamics, and monitor how they evolved over time, we used a trajectory analysis. For this, we used spike counts from 50 ms time bins across all recorded neurons as a high dimensional vector. These vectors were projected onto three dimensions that captured a large fraction of the variance (**Fig. 1D**; Principal Component Analysis or PCA; n = 66 PNs; ∼ 50% variance captured). As expected, neural response vectors during ON periods (orange trajectories; 4 s during the odor presentation) were distinct from response vectors during OFF periods (blue trajectories; 4 s from the termination of odorant) for the duration of the 4 sec OFF response window.

Next, to quantitatively examine the dissimilarity between ensemble ON and OFF responses, we applied a correlation analysis using the high-dimensional response vectors. For this we calculated pairwise correlations between all PN response vectors observed at different time points as a correlation matrix (**Fig. 1E**). Each row or column of the correlation matrix indicates the similarity between an ensemble response vector at a specific time point with all other PN response vectors observed over time. Consistent with the observed neuronal activity and PCA trajectory analysis, neural responses over time within the ON period (for the duration of the odor presentation) were highly correlated amongst one another (ON-ON). Similarly, OFF responses were highly correlated with one another (OFF-OFF). However, the cross-correlation between any ON response timepoint with any OFF response timepoint (ON-OFF) was greatly reduced compared to intra-period correlations. These results show that ON and OFF neural response neural responses were quantitatively distinct from one another.

Finally, we examined whether the distributions of angles between ON and OFF response vectors (a measure of dissimilarity) were consistent across a panel of diverse odors. For each odor, we calculated the angle between the high-dimensional PN response vectors for timepoints during the odor presentation and after the odor presentation. Vector comparisons fell into three groups: (1) comparing vectors from the ON period to other vectors from the ON period (ON vs ON; 80 vectors, 3160 pairwise comparisons), (2) comparing vectors from the ON period to vectors from the OFF period (80 ON vs 80 OFF vectors; 6400 pairwise comparisons) and (3) comparing vectors from the OFF period to other vectors from the OFF period (OFF vs OFF; 80 vectors, 3160 pairwise comparisons). The distributions of angles were consistent across odors (**Fig. 1F**, **Fig. S1**). In contrast, angles between random vectors of the same dimensionality clustered with means and medians around 90° (gray distributions). To allow quantitative comparisons across odors, we computed the median angular separation for each odor per category (**Fig. 1G**). For every odor the median angle between the ON and OFF responses was more than 90 degrees (mean = 106.7° +/- 0.84 SEM, n = 11 odors), reflecting the negative correlation between ON and OFF responses (**Fig. 1D**). These data are consistent with our prior results^27^, and generalize our previous findings by showing that distinct neural ensembles are activated during and after presentations of a wide panel of odors.

Distinct odor ON and OFF responses have also been described in the mouse OB^24^, but the relationship between ON and OFF responses is unknown. To test whether the differences between ON and OFF population response dynamics in the locust are also observed in the mouse, we performed a similar analysis of odor evoked responses recorded from the mouse OB. For this, we imaged glomerular-layer mitral and tufted cell dendrites in Thy1-GCaMP6f mice using a two-photon random access mesoscope^28^ during passive odor presentations (**Fig. 2A, Fig. S2**). With this approach, we monitored calcium signals from hundreds of glomeruli (median: 215 glomeruli per imaging session) across the dorsal surface of bilateral OBs with high spatial and temporal resolution (**Fig. 2B-C**).

**Fig. 2:**
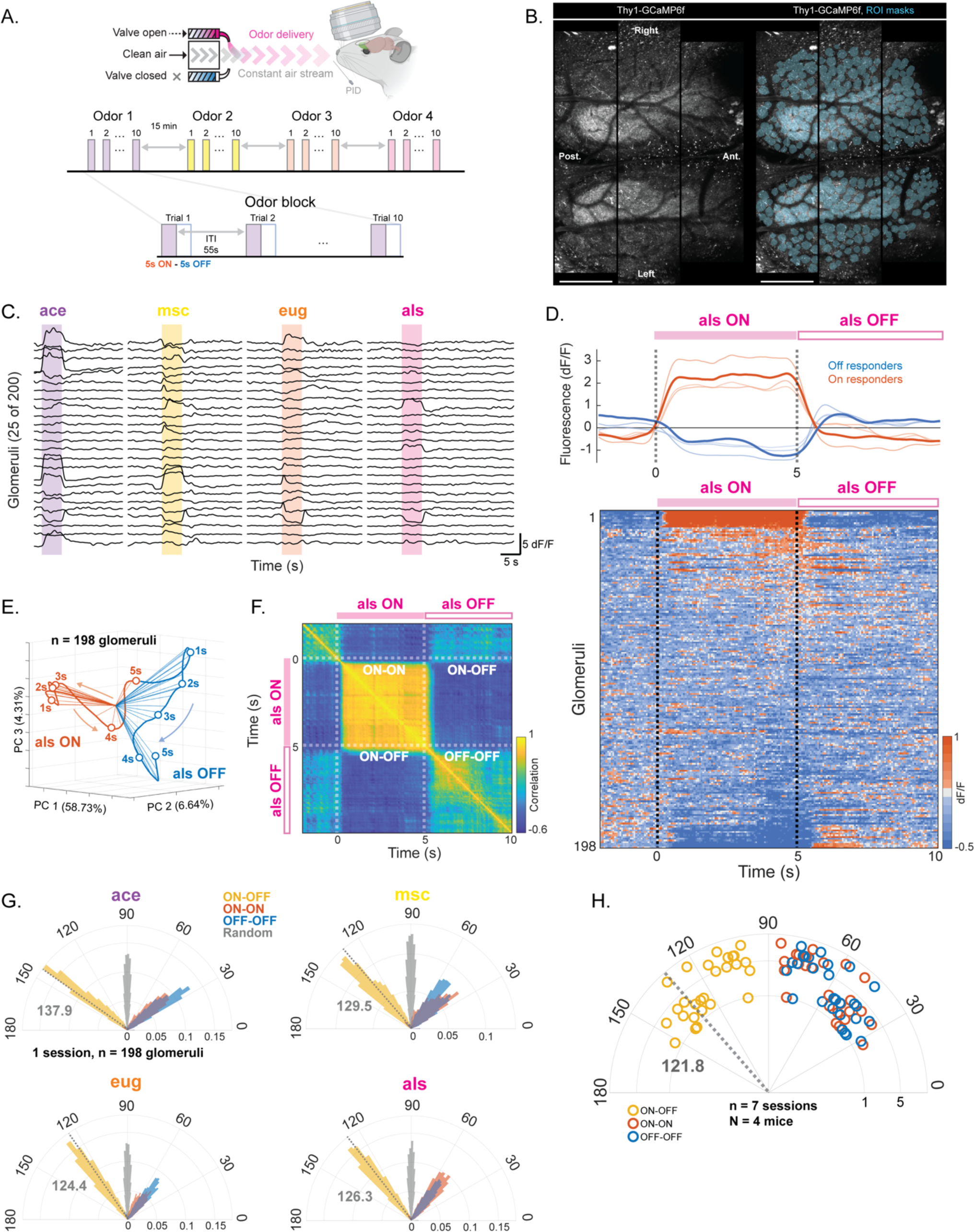
Odor ON and OFF responses in the mouse olfactory bulb are anticorrelated. **A.** Schematic of the experimental design. (top) Neuronal activity was observed with mesoscale two-photon calcium imaging through chronic cranial windows over the mouse OB. (bottom) Odors were presented for five seconds, separated by 55 second intertrial intervals, in blocks of ten trials. 15-minute periods of no odor stimulation separated blocks of different odors. **B.** Representative trial-averaged image of GCaMP fluorescence in the olfactory bulb of a Thy1-GCaMP6f mouse (left) shown with ROIs overlaid on glomeruli (right) from bilateral olfactory bulbs. Scale bar is 500 um. **C.** Fluorescence traces (dF/F) from 25 representative glomeruli showing responses to acetophenone (ace, purple), methyl salicylate (msc, yellow), eugenol (eug, orange), and allyl sulfide (als, pink). **D.** (top) Average fluorescence (dF/F) trace from subset of ON (orange) and OFF (blue) responsive glomeruli. (bottom) Heatmap showing trial-averaged dF/F traces from all glomeruli in one mouse in response to allyl sulfide (als, pink bar) and after odor offset. **E.** PCA trajectories of trial averaged glomerular population response during allyl sulfide presentation (als ON, orange) and after allyl sulfide presentation (als OFF, blue). **F.** Representative correlation matrix summarizing the similarity between trial-averaged glomerular responses evoked by allyl sulfide (als, pink bar) during ON and OFF time periods. **G.** Distributions of angular distance between pairs of glomerular response vectors for all four odors presented (ace, msc, eug, als). Angular separations are calculated between vectors within the ON (orange) and OFF (blue) periods and between any ON response vector and any OFF response vector (yellow). Distribution of angular separation between random vectors (grey) are shown as a control. Median values of ON vs OFF angular separation distributions indicated with dashed lines and grey text. **H.** Summary of the median angular distances for 4 odors (n = 7 imaging sessions from 4 mice) showing ON vs ON period median angles (orange), OFF vs OFF period median angles (blue) and ON vs OFF period median angles (yellow). The mean of the ON vs OFF angles is shown as the dashed grey line and text. Radii indicate different imaging sessions (n = 7 imaging sessions from = 4 mice).

We found that, like locust PNs, distinct sub-populations of mitral and tufted cells uniquely responded to odor onset or odor offset (**Fig. 2D**). These data are consistent with previous studies reporting that neural responses in the mouse olfactory bulb continue after termination of a stimulus^24^. Our data further revealed that glomeruli responding to odors during odor presentation tended to be suppressed following odor offset, and that glomeruli responding most strongly to odor offset were suppressed during the odor presentation (i.e., ON and OFF responses were mutually exclusive (**Fig. 2D**, traces). Dimensionality reduction by principal components showed that the temporal trajectories of both ON and OFF responses to single odor presentations were distinct (**Fig. 2E**). Intriguingly, the pairwise correlation of glomerular ON and OFF response vectors revealed that odor ON and OFF responses were not just distinct, but were anticorrelated. In contrast, angular separation between scrambled vectors clustered around 90° (**Fig. 2F**). Thus, odor ON and OFF responses in the mouse OB displayed a remarkably similar relationship to that observed in the locust (**Fig. 2E, F** *vs.* **Fig. 1D, E**). In high-dimensional feature space, the angle between ON and OFF trajectories was greater than 90° for each odor presented (mean = 121.8° +/- 2.98 SEM, n = 4 odors, 7 sessions) (**Fig. 2G, H**), revealing that, as in the locust, odor ON and OFF responses in mouse OB glomeruli are inverted relative to each other, and not merely independent (orthogonal) responses. Together these data suggest that odor-evoked aftereffects in the mouse OB and locust AL form an inverted odor afterimage. It is worth noting that the anticorrelation between ON and OFF responses imaged from dendrites in the mouse OB is larger than that observed from locust PN recordings. This is unlikely to be the result of differences in the temporal dynamics of our recording modalities given that, (1) we would expect to observe sharper transitions from ON to OFF and a stronger anticorrelation using the method with higher temporal resolution, and (2) in locust experiments, action potentials are pooled into 50ms bins to calculate firing rates, making the temporal resolutions of the two recoding modalities comparable. More likely, the stronger anticorrelation observed in the mouse OB results from imaging populations of glomeruli simultaneously in individual trials while pooling dendrites from correlated sister cells innervating the same glomeruli, thus reducing some neuron-to-neuron and trial to trial variability.

### ON and OFF responses occupy distinct regions of feature space

Next, we examined the separability of population odor responses by odor identity during ON and OFF time periods. Dimensionality reduction revealed that each ON response formed a distinct trajectory, indicating that unique populations of PNs were activated during each odor’s ON period (**Fig. S3A**). Similarly, OFF responses formed distinct trajectories, indicating that they were also odor-specific (**Fig. S3B**). We then performed principal component analysis using neural responses during both ON and OFF periods and found that, across odors, ON and OFF periods occupied different regions of PCA space. This suggests that ON responses for different odors were markedly more similar to each other than they were to OFF responses across odors (**Fig. 3A**). To quantitatively validate the observations from PCA analysis, we used high-dimensional response vectors to calculate angular separation between the ON response elicited by a reference odor with the ON and OFF responses elicited by all odors in the odor panel (**Fig. 3B**). We found that separations between the ON response of the reference odor and the ON responses to all other odors were smaller than the separations between the ON response for the reference odor and the OFF response to all odors (including the reference). While this finding was consistent for most reference odors, the difference in angular separation was significant for 5 out of 11 odors in the panel (**Fig. S4A**).

**Fig. 3:**
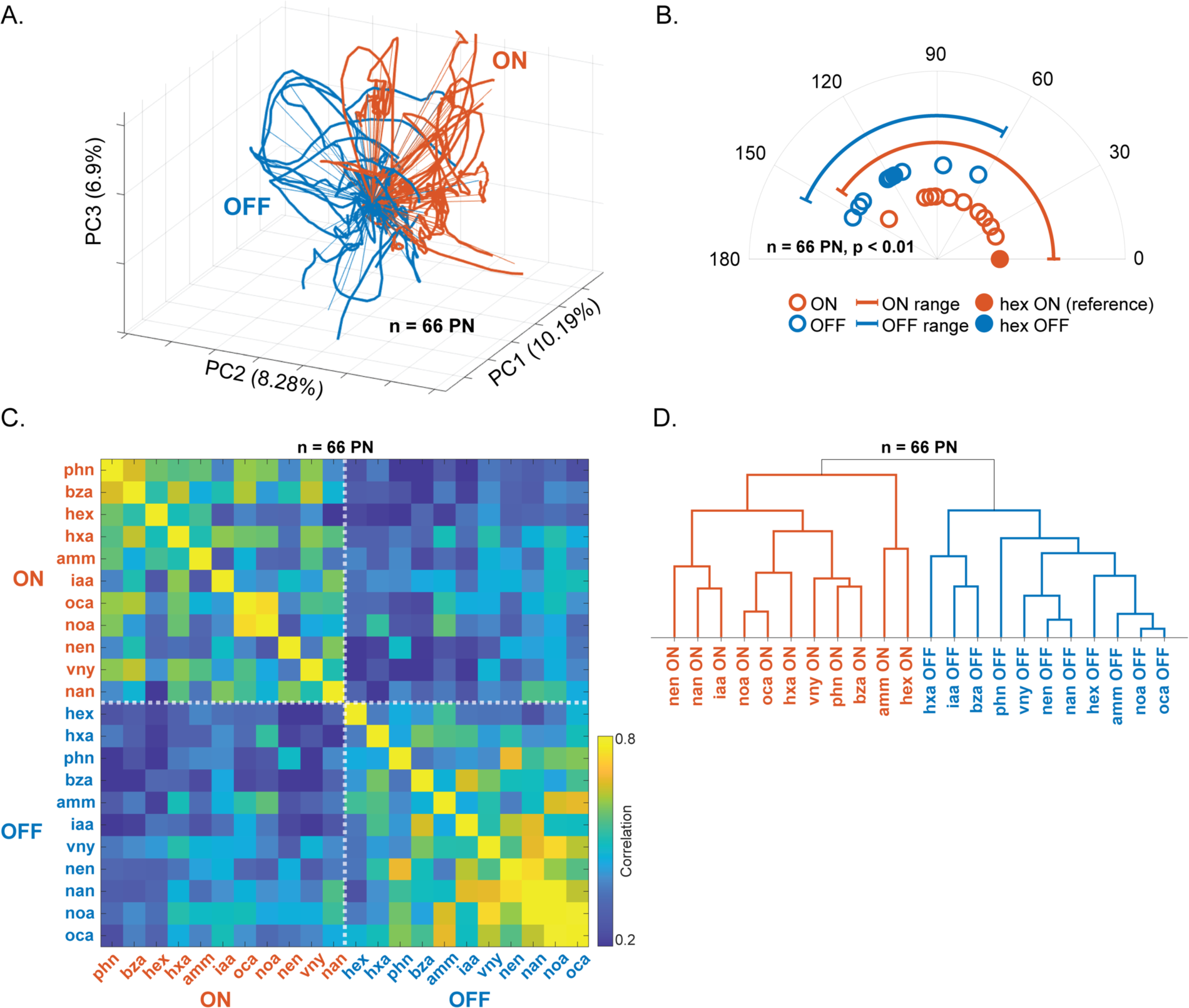
ON and OFF responses in locust antenna lobe contain distinct odor specific information. **A.** PCA trajectories of odor-evoked ON (orange) and OFF (blue) responses to all 11 odors in the panel are shown. **B.** Polar plot of angular distances between the hex ON response vector (reference, solid orange circle; averaged over 4 seconds odor presentation time window) and the ON response vectors of the 10 other odors (orange open circles). Angular distances between the hex ON response vector and the OFF responses of all odors (blue open circles) including hex OFF response (solid blue circle) are shown. Arcs (thick lines) show the range of angular distances observed for ON-ON angular distances and compares the same with ON-OFF angular distances. **C.** Correlation of trial-averaged PN ON and OFF responses to all eleven odorants in the panel. **D.** Hierarchical clustering of ensemble PN ON and OFF responses are shown.

To further evaluate this result, we next calculated the correlations between ON and OFF responses of all odors in the panel (**Fig. 3C**). As expected, intra-period correlations (ON vs. ON and OFF vs. OFF) were relatively high. In comparison, the inter-period correlations (ON vs. OFF) were much lower. Hierarchical clustering of high dimensional ON and OFF responses revealed that PN responses during ON and OFF periods formed distinct response clusters (**Fig. 3D**).

Consistent with the locust results, in mice, we found that both ON and OFF responses were odor-specific (**Fig. S3C, D**), and that ON responses occupied a distinct region of PCA space compared to OFF responses (**Fig. 4A**). As in locusts, angular separations between a reference odor and all other odor ON responses were smaller than angular separations between the reference ON response and all odor OFF responses (**Fig. 4B**). This result was true for all odors in the panel (**Fig. S4B**). Correlating glomerular response vectors across ON and OFF periods of different odor presentations further demonstrated that ON responses were distinct from OFF responses in general (**Fig. 4C**). Distributions of correlations across odors revealed systematically lower correlations for inter-period compared to intra-period correlations in both mouse and locust (**Fig. S5**). Supporting this, hierarchical clustering of mouse odor responses cleanly separated ON and OFF responses (**Fig. 4D**), showing that, as in the locust AL, ON and OFF responses in the mouse OB represent distinct patterns of neuronal activity.

**Fig. 4:**
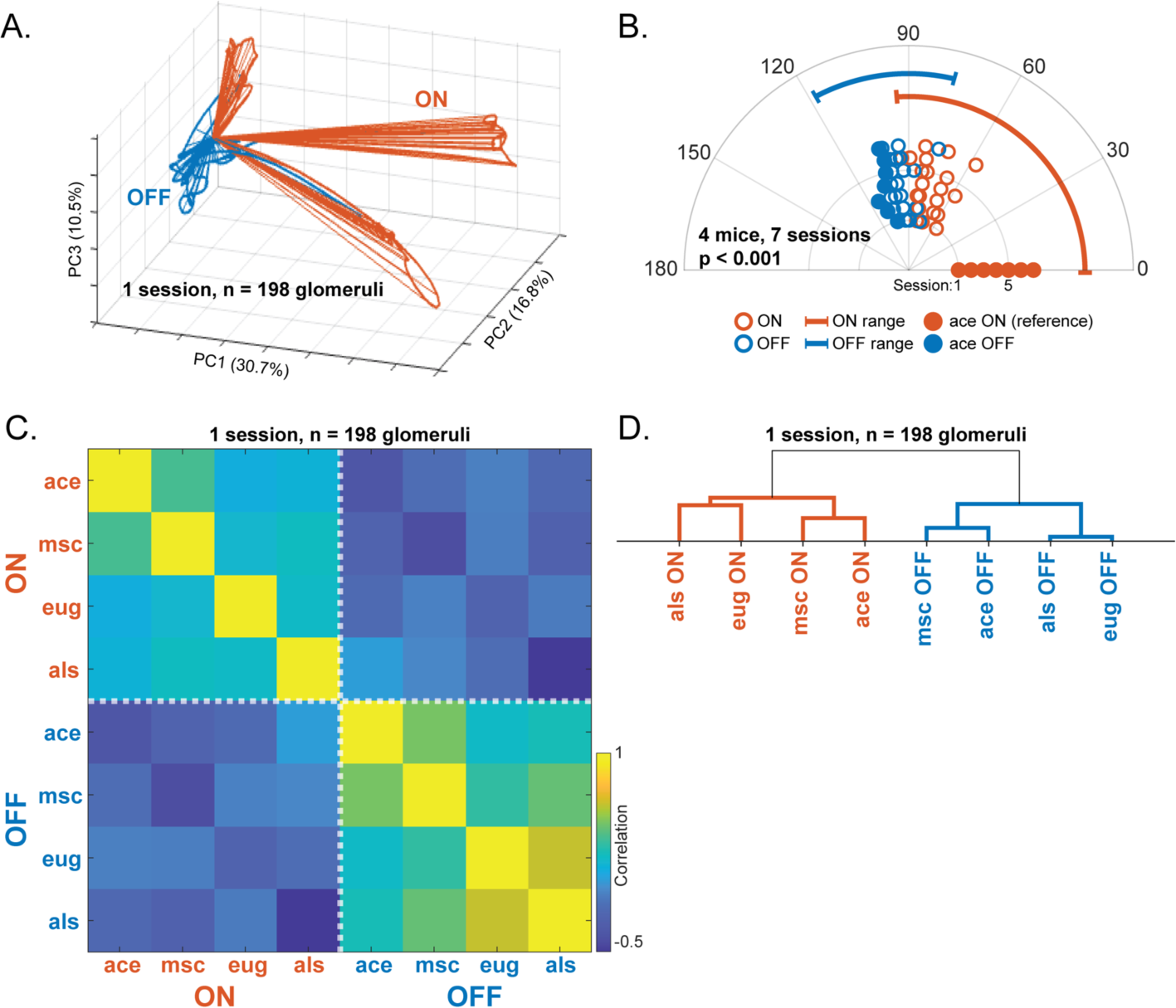
ON and OFF responses in the mouse olfactory bulb contain distinct odor specific information. **A.** PCA trajectories of odor-evoked ON (orange) and OFF (blue) responses to four odors: ace, msc, eug and als. **B.** Polar plot showing the angular distance between ace ON response vector (reference, solid orange circles), ON response vectors of the 3 other odors (open orange circles), ace OFF response vector (solid blue circle), or OFF response vector of the three other odors (open blue circles). Radii indicate responses from different imaging sessions (n = 7 imaging sessions from 4 mice). Arced lines show the range of angular distances between ON responses (orange). **C.** Correlation matrix summarizing the similarity between glomerular ON and OFF responses evoked by four odors. **D.** Hierarchical clustering of glomerular ON and OFF response vectors from a representative imaging session.

### History-dependent suppression enhanced between sequentially presented odors

Given that odors can be recognized within hundreds of milliseconds of stimulus onset^29^, how can OFF responses, which begin after a stimulus has been terminated, contribute to odor processing? One possibility is that neural activity after stimulus offset could interfere with subsequent odor responses. To test this hypothesis, we presented locusts with pairs of odors in rapid succession, such that the OFF response of the first odor would overlap with the ON response of the second odor in the sequence (**Fig. 5A**). In locusts, we found that PNs that selectively responded to only one odor in the sequence maintained those responses whether the odors were encountered solitarily or in a sequence (i.e., unique responses were unperturbed; **Fig. 5A**, PN 1 and 2). Whereas PNs that responded to both odors individually, responded only to the first odor when the two odors were presented in sequence (**Fig. 5A**, PN 3). The response to the second odor was suppressed (i.e., overlap reduction).

**Fig. 5:**
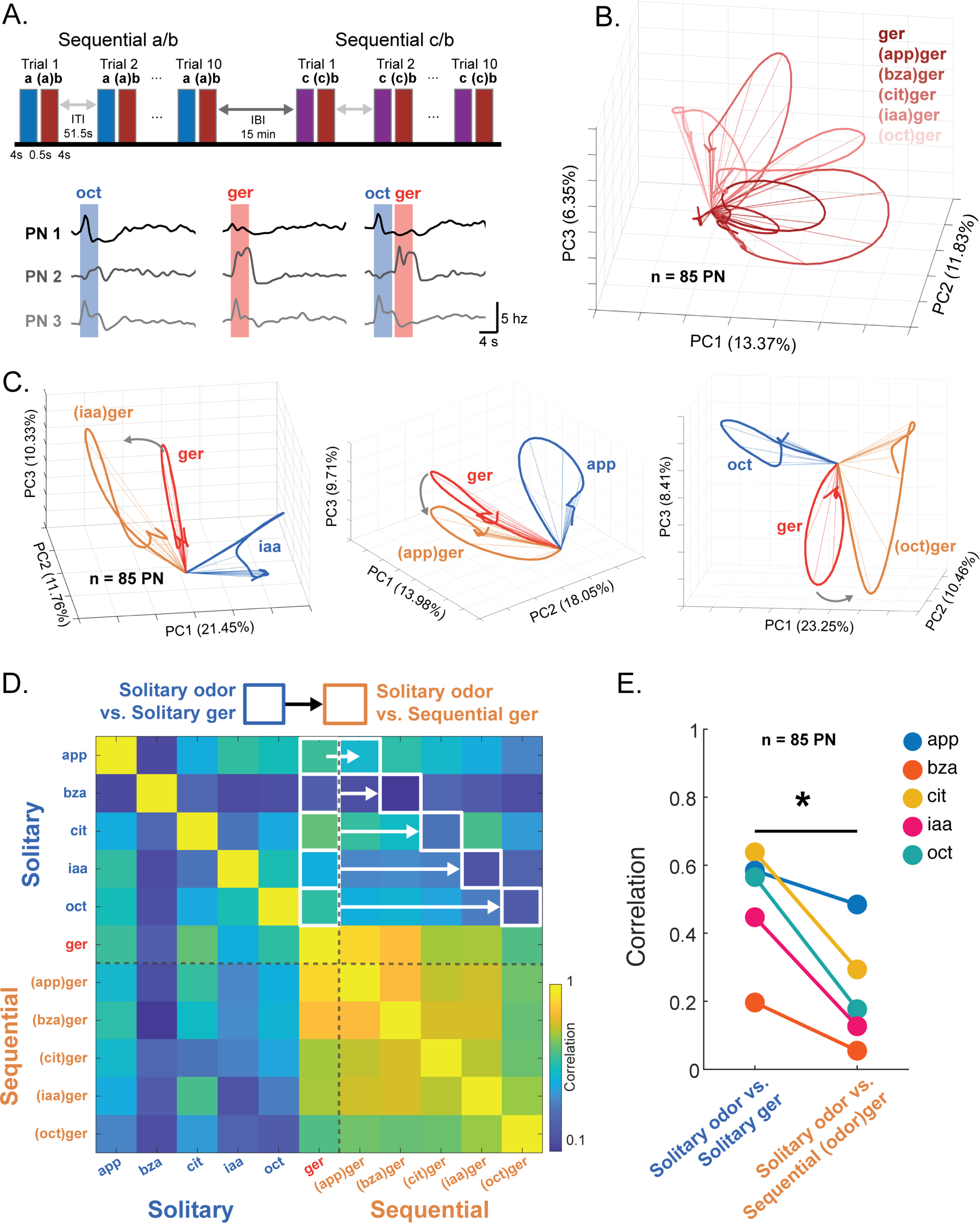
Contrast is enhanced in the locust antenna lobe between odors presented sequentially. **A.** (top) Experimental design for sequential blocks. Odors presented include geraniol (ger), isoamyl acetate (iaa), benzaldehyde (bza), citral (cit), octanol (oct), and apple (app). 4 second pulses of Odor A were presented with 4 second pulses of Odor B presented 0.5 second afterwards during the OFF period of Odor A. (bottom) Example responses of two projection neurons (PN) to oct (left) and ger (middle) presented individually and to ger presented in sequence with oct (right). **B.** PCA trajectory of ger ON responses. Traces represent average PN response during the ON period to ger alone (ger), or ger with different odors that presented immediately prior (sequential: (app)ger, (bza)ger, (cit)ger, (iaa)ger, (oct)ger). **C.** (left) PCA trajectory ON responses to solitary iaa (blue), solitary ger (red), and ger sequentially after iaa ((iaa)ger, orange). (middle) PCA trajectory ON responses to solitary app (blue), solitary ger (red), and ger sequentially after app ((app)ger, orange). (right) PCA trajectory ON responses to solitary oct (blue), solitary ger (red), and ger sequentially after oct ((oct)ger, orange). **D.** Correlation of average PN ON responses to solitary odors (app, bza, cit, iaa, oct, ger) and to ger presented in sequence after other odors ((app)ger, (bza)ger, (cit)ger, (iaa)ger, (oct)ger). White boxes and arrows indicate which correlation values are used for the quantitative comparison in (E). **E.** Correlation values of PN population responses to solitary ger vs PN population responses to other solitary odors (left boxes in D) compared to correlation values of PN responses to solitary odors vs PN responses to ger presented sequentially after the other odors (right boxes in D, one-tailed t test, p < 0.05, n = 5 odor pairs).

Given that individual PN responses evoked by an odorant depend on previously encountered stimuli, we expected AL ensemble activity to vary systematically as a function of prior odor exposure. To test this, we first visualized PN neural responses evoked by a single odorant (geraniol, ger) encountered by itself (“solitary”) or as the second odor in a two-odor sequence (“sequential”) (**Fig. 5B**). As expected, presentations of geraniol after other odors produced response trajectories that relied on the prior odor in the sequence. Comparing response trajectories evoked during the solitary and sequential geraniol presentations, we found that geraniol responses during the sequential encounters became more distinct (greater angular separation) with respect to the first odor in that sequence (blue trajectories in **Fig. 5C**). This was observed for each odor sequence tested. Consistent with PCA trajectory analysis, we found that the correlation between high-dimensional odor response vectors was lower when the odors were sequentially presented, compared to when the odors were presented in isolation (**Fig. 5D, E**). Together, these data support the hypothesis that inverted OFF responses among locust PNs allow AL circuits to enhance response contrast between sequentially encountered odorants.

We next questioned if the observed OFF responses serve a similar role in enhancing contrast between odor representations in the mouse OB. To test this directly, we presented mice with either solitary odors, or odors in sequence (**Fig. 6A**). As observed for locust PNs (**Fig. 5**), overlapping subsets of glomeruli responded to individual odors when presented solitarily. However, when those same odors were presented sequentially, the glomeruli that responded to the first odor were suppressed during the presentation of the second odor (**Fig. 6A**, traces). As in locust, suppressing overlapping glomerular activity caused response trajectories for the same odor to vary depending on stimulus history (**Fig. 6B**, **Fig. S6A**), and ensemble trajectories of sequentially presented odors reliably deflected away from the trajectory of a previously presented odor (**Fig. 6C**, **Fig. S6C**). This indicated that responses to odors became more distinct (i.e., decorrelated) during the sequential encounters. To quantify this observed history-dependent contrast enhancement, we computed correlations between high-dimensional odor response vectors when odors were presented solitarily, and compared these values with the correlations between responses when the same odors were presented sequentially (**Fig. 6D**, **Fig. S6B**). The qualitative trajectory analyses, together with the high-dimensional correlation analyses showed that when odors were presented sequentially, responses in the mouse OB became less similar (**Fig. 6E**). Together, these results suggest that contrast enhancement by inverted odor OFF responses represent a computation of early olfactory circuitry that is conserved across locusts and mice.

**Fig. 6:**
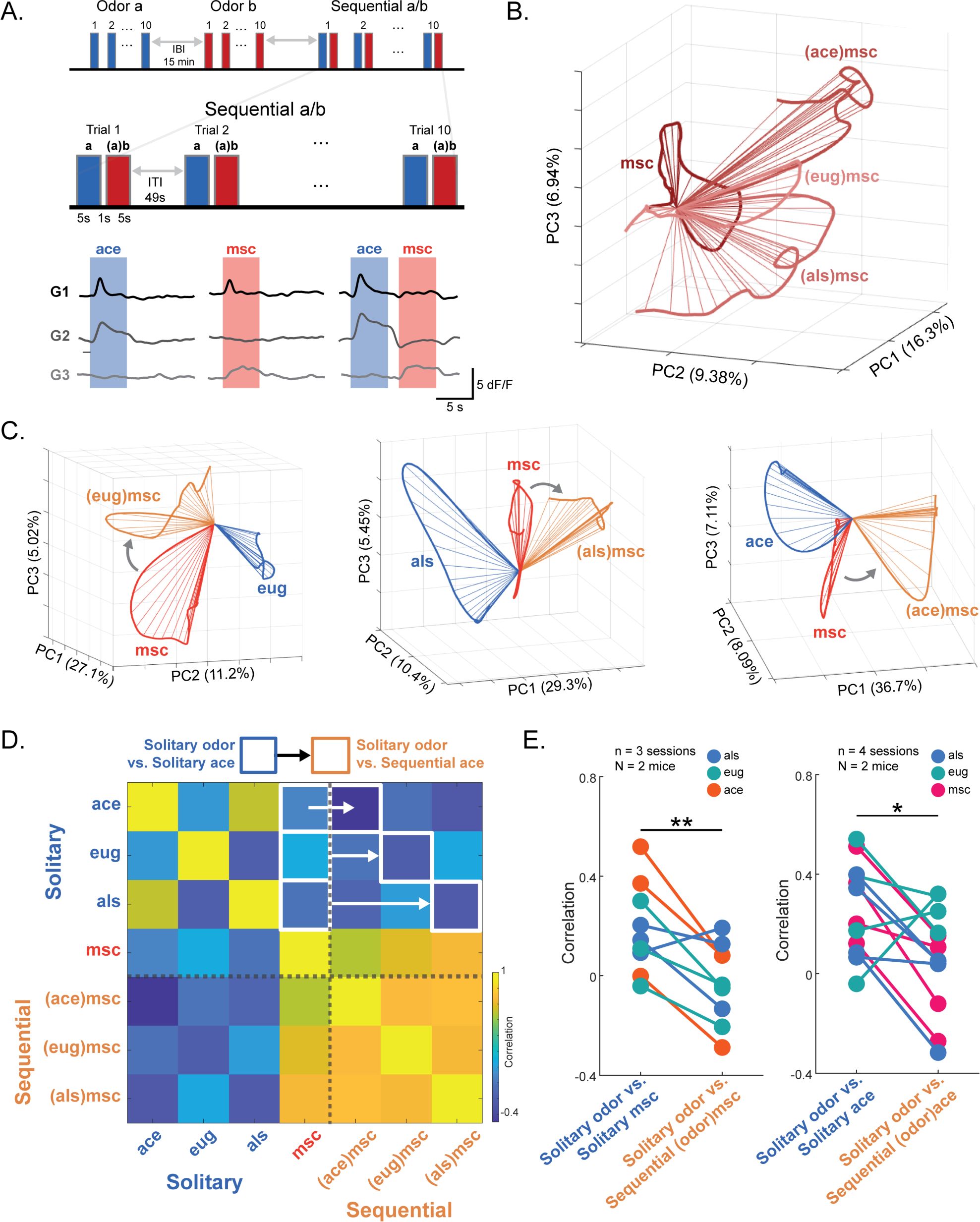
Contrast is enhanced in the mouse olfactory bulb between odors presented sequentially. **A.** (top) Schematic of odor presentations showing solitary presentations of single odors and sequential presentations of two odors with a one second interval between paired odors in a trial. Odors presented include acetophenone (ace), methyl salicylate (msc), eugenol (eug), and allyl sulfide (als). 5 second pulses of Odor A were presented with 5 second pulses of Odor B presented 1 second afterwards during the OFF period of Odor A. (bottom) Example responses of three glomeruli (G) to ace (left) and msc (middle) presented individually and to msc presented one second after ace (right). **B.** Representative PCA trajectories showing the response to a solitary presentation of msc (dark red) and responses to msc sequentially presented after ace ((ace)msc), eug ((eug)msc), and als ((als)msc, lighter reds). **C.** Representative PCA trajectories showing responses to solitary and sequential presentations of msc. (left) solitary eug (blue), solitary msc (red), and msc sequentially presented after eug ((eug)msc, orange). (middle) solitary als (blue), solitary msc (red), and msc sequentially presented after als ((als)msc, orange). (right) solitary ace (blue), solitary msc (red), and msc sequentially presented after ace ((ace)msc, orange). **D.** Correlation of glomerular ON responses across solitary presentations of eug, ace, msc, and als and sequential presentations of msc following ace ((ace)msc), eug ((eug)msc), and als ((als)msc). White boxes and arrows indicate correlation values used for the quantitative comparison in (E, left). **E.** (left) Correlation values between solitary presentations of msc and other odors (left boxes in D) compared to correlation values between sequential msc and odors across imaging sessions (right boxes in D, n = 3 imaging sessions from 2 mice; one-tailed t test, p < 0.01) (right). Correlation values between solitary presentations of ace and the three other odors (see also Figure S5) compared to correlation values between sequential ace and odors across imaging sessions (n = 4 imaging sessions from 2 mice; one-tailed t test, p < 0.05).

### Odor OFF responses persist long after odor offset

OFF responses in locust AL and mouse OB represent the persistence of odor-specific information after the termination of the stimulus, and therefore may provide a neural basis for a form of short-term memory. This raised the question of how long, and in what form, odor information persists in olfactory neuronal networks. To examine this, we first observed how post-stimulus network activity in the AL and OB relates to odor-evoked responses. As shown in prior analyses, distinct subsets of locust AL PNs (**Fig. 1**) and mouse OB glomeruli (**Fig. 2**) respond to odor onset (ON responders) and odor offset (OFF responders). We found that the activity of ON responders continued to be suppressed, and the activity of OFF responders was elevated for several seconds after stimulus offset in both locust AL (**Fig. 7A**, **S7A**) and mouse OB (**Fig. 7B**). Further, neuronal activity in the AL and OB maintained negative correlations with the odor-evoked ON responses for several tens of seconds following stimulus termination (Locusts – **Fig. 7C, S7B**, Mouse – **Fig. 7D, S7C**). This result suggests that although the overall neural activity after stimulus termination returns rapidly to pre-stimulus levels, network activity remains negatively correlated or inverted with respect to the odor-evoked response.

**Fig. 7:**
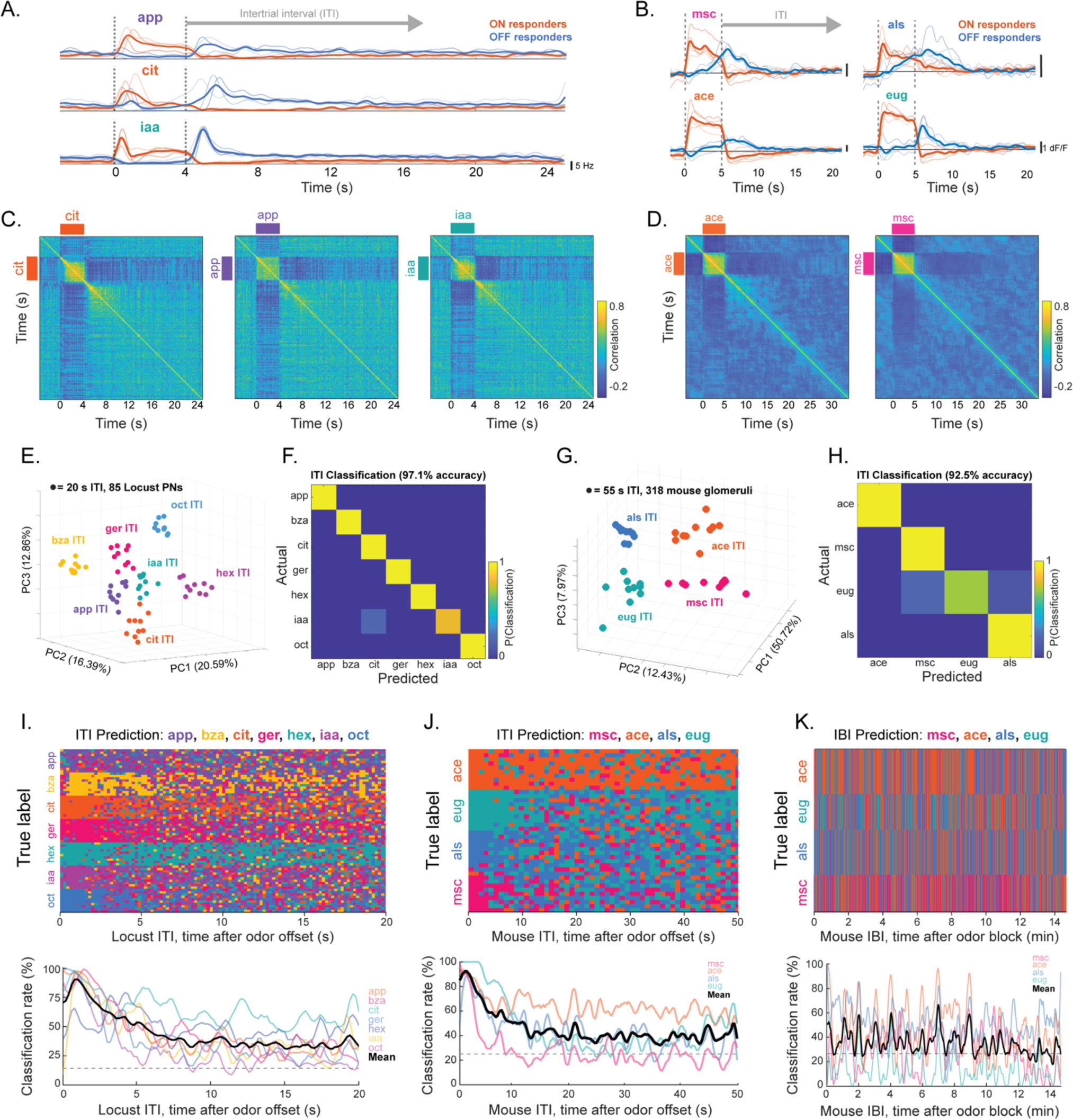
OFF responses persist long after odor presentations in mouse OB and locust AL. **A.** Trial averaged PSTH traces from representative ON (thin orange) and OFF responsive (thin blue) PNs responsive to apple (app), citral (cit), or isoamyl acetate (iaa). Thick lines are means. **B.** Trial averaged dF/F traces from representative ON (thin orange) and OFF (thin blue) responsive glomeruli in mouse OB showing pre odor, odor-evoked, and post-odor activity for methyl salicylate (msc), acetophenone (ace), allyl sulfide (als), and eugenol (eug) with mean dF/F of ON (thick orange) and OFF responders (thick blue). **C.** Representative temporal correlations of PN responses to app, cit, and iaa. **D.** Representative temporal correlations of OB glomerular responses to ace and msc. **E.** Dimensionality-reduced ensemble PN activity during 20 seconds after odor offset during intertrial intervals (ITI) between presentations of the same odor color coded by odor. **F.** Results from a K-NN classification analysis of PN activity using angular separation of full-dimensional data as the distance metric. **G.** Dimensionality-reduced mouse OB glomerular activity during the 55 second ITIs between presentations of the same odor within a block of 10 trials color coded by odor. **H.** Results from a K-NN classification analysis of the glomerular activity (k = 20) during the 55 second ITI period, based on angular distance of full-dimensional data. **I.** (top) Time-bin-by-time-bin k-NN classifier (k = 10; correlation distance; locusts PN activities) predictions for the post stimulus ITI time bins color coded by odor. (bottom) Quantification of classifier accuracy during the ITI time window for each odor (colored transparent lines) and averaged across odors (black line). Grey dashed line represents chance level. **J.** (top) Time-bin-by-time-bin k-NN classifier (*k* = 10; correlation distance; mouse glomerular activities) predictions of the post stimulus ITI time color coded by odor. (bottom) Quantification of classifier accuracy during the ITI time window (colored transparent lines) and averaged across odors (black line) are shown with chance as dashed grey line. **K.** (top) Time-bin-by-time-bin k-NN classifier (*k* = 50; mouse glomerular activities) predictions of the IBI time color coded by odor. **J.** (bottom) Quantification of classifier accuracy for the IBI following each odor (colored transparent lines) and averaged across odors (black line) with chance (grey dashed line).

The persistence of stimulus-specific activity patterns (an odor afterimage) in the locust AL and mouse OB indicated that odor identity could be inferred based on the post-stimulus ensemble activity, well after stimulus offset. To test this, we used PCA analysis to visualize post-stimulus neural activity and found that activity during inter-trial intervals (ITIs) clustered robustly by odor identity (**Fig. 7E**). A k-nearest neighbor classifier, using angular distances of the full-dimensional ITI data, robustly discriminated odors (**Fig. 7F**). This confirmed the interpretation from the PCA analysis and showed that ITI activity was odor-specific. We observed a similar clustering of 55-second ITI activity from the mouse OB (**Fig. 7G**), which contributed to highly accurate classification of odors from ITI glomerular responses (**Fig. 7H**). Additionally, we repeated this classification analysis using one-minute bins of glomerular activity taken from a 15-minute window between blocks of different odors (inter-block interval, IBI, **Fig. S7D**). Our results indicate that during the IBI, network activity ceased to contain odor-specific information. We observed no clustering of binned responses taken from the IBI, and classification accuracy was at roughly chance levels (**Fig. S7E**).

Complementing the time-averaged classification analysis spanning the entire ITI period, we also performed a classification analysis over time. As expected, classifier performance rapidly increased at odor onset and remained elevated for the duration of the odor presentation in both locusts and mice (**Fig. S8A, B**). After odor offset, we found that among locust PNs, odor information decayed gradually across the ITI. Classifier accuracy decreased to ∼40% across odors throughout the 20-second post stimulus ITI, but notably, for most odors, accuracy did not fall to chance levels during the entire ITI period (**Fig.7I**). In the mouse OB, the persistence of odor information was also striking. We found that classifier accuracy remained high for the duration of the 55-second ITI (**Fig.7J**). Although classifier accuracy was variable across odors, PID recordings confirmed that, for all odors, concentrations near the nares decayed rapidly after odor offset **(Fig. S2).** Shuffling odor labels results in chance-level classifier performance throughout the ITI (**Fig. S8C**). Additionally, classifier accuracy was greatly decreased and was near chance levels for all odors during the 15-minute IBI (**Fig.7K**).

To determine whether odor information persisted in the ITI after single presentations of an odor (as opposed to repeated presentations of the same odor) we performed a similar set of experiments in mice, where eleven odors were presented in a random order and classifier performance was measured throughout the ITI. In these experiments, odors were presented for 1 second, with 19 second ITIs and no block structure. Classifier accuracy across the ITI was variable across odors (**Fig. S8D**), but, on average, declined to chance after several seconds (**Fig. S8E**). This is in contrast to the repeated odor presentation condition where accuracy remained above chance for the duration of the 55 second ITI (**Fig. 7J**). This is likely due to the difference in odor presentation structure between the two experiments, with accumulating hysteresis contributing to the extreme persistence of odor information that we see in the repeated odor presentation condition. Collectively, these findings indicate that AL PNs and OB glomeruli maintain information about an odor stimulus well after its termination.

## DISCUSSION

The goal of this study was to identify common computational principles governing early stages of olfactory processing across species. For this purpose, we used calcium imaging and electrophysiological recordings to compare the dynamics of odor-evoked responses in the mouse OB and the locust AL, respectively. A strength of this comparative approach is that we leveraged unique strengths of each model and recoding modality. In the mouse OB, genetically targeted calcium indicators and two-photon imaging allowed us to isolate and image postsynaptic dendritic responses in the glomerular layer. Due to the rigidly stereotyped anatomy of mouse OB circuits, these dendritic responses were easily grouped into glomerular units comprised of highly correlated sister mitral cells. Thus, calcium imaging from dendrites in the glomerular layer of the mouse OB provided a way to pool signals from functionally-related mitral cell cohorts while sampling a broad population of odor responsive neurons postsynaptic to OSNs. Furthermore, though calcium indicators only approximate neuronal action potential firing, in the current study we were specifically interested in dendritic responses upstream of cell bodies regardless of whether this activity ultimately led to action potentials in some or all of the correlated sister cells. At the same time, single cell electrophysiology in the locust provided a complimentary window into odor response dynamics by providing high-temporal resolution information about action potential firing in individual cells. While just a few units were simultaneously recoded in locust AL, population odor responses were collected across multiple experiments. As such, multi-unit recordings in locusts allowed us to record from AL PNs without genetic targeting and quantify AL PN output in the form of action potential firing downstream of complex sensory neuron input, local circuit modulation, or intrinsic changes that might occur at the cell body.

Despite differences in the population breadth and kinetics of our electrophysiological and imaging modalities, as well as the structural differences in locust and mouse olfactory systems, our comparative approach revealed surprisingly conserved odor response dynamics across species. First, we found that responses to odor onset (ON responses) and odor offset (OFF responses) are distinct and anticorrelated. We then demonstrated that OFF responses interfere with subsequent odor presentations and enhance contrast between sequentially presented odors in both mouse and locust. Finally, we found that inverted OFF responses persist and maintain short-term odor memories over several seconds. Ultimately, our comparative approach reveals functional consequences of long-lasting, temporally-complex odor-evoked activity that generalize across species, even if the quantitative details of these effects are different, and likely influenced by recording modality, model, and experimental design. Taken together, our findings support the notion that odor response dynamics are conserved across systems to serve core functions of early olfactory circuits in both the locust AL and mouse OB.

Persistently altered neural dynamics and distortions in perception after an encounter with a stimulus – “aftereffects” – are a ubiquitous feature of sensory systems across species, and have well described perceptual and behavioral consequences in visual, auditory, and somatosensory systems. Importantly, the perceptual distortions of aftereffects reveal computations performed by sensory circuits that are critical for stimulus detection and discrimination. Aftereffects across sensory modalities and species correspond to changes in neuronal activity with stimulus offset – either through neuronal adaptation that occurs during stimulus presentation, or through distinct stimulus offset-evoked responses^30–35^. In olfaction, aftereffects have been described in the form of distinct neural responses to the offset of odors in worms, insects, fish, and mammals^24,27,36–38^. In the mammalian olfactory system, previous studies have shown that odor-specific information briefly persists in the OB after stimulus offset^24^. In contrast, we found that patterns of glomerular activity persisted for several seconds after individual odor presentations (**Fig. S8D, E**), and for the entire duration of the ITI when odor presentations were repeated (**Fig. 7J**), allowing odor identification for up to a minute after presentations. Our results build on previous studies to suggest that odor memories, in the form of ‘after images’, are longer lasting than previously appreciated. Notably, the anticorrelation of OFF responses in mice and locusts arises from the suppression of ON responding neurons and glomeruli during the OFF period, and the suppression of OFF responding neurons and glomeruli during the ON period. At the same time, distinct sets of neurons were activated during ON and OFF periods. Whether this is a generic strategy for retaining stimulus specific information in neuronal networks over time, how much stimulus specific information is encoded in the suppression of ON responders vs the activation of OFF responders, and whether complementarity in activated sets of neurons also forms the basis for other adaptive computations remains to be seen. Importantly, the current work reveals a specific form of persistent population neural activity, anticorrelated to odor-evoked responses, which allows information about recently encountered odors to influence subsequent odor responses.

Given the distinct nature of odor OFF responses, and the surprising finding that odor specific activity persists long after odor offset, an intriguing question is what cellular and circuit mechanisms drive these effects, and are they fixed or flexible? In locusts, it has been shown that (1) odor offset-evoked activity is distinct from odor onset-evoked activity, and (2) that OFF responses are anticorrelated with ON responses through a combination of cell-intrinsic properties and local GABAergic circuit interactions^27^. Part of this inverted OFF response relies on adaptive changes in spike-thresholds of individual PNs, such that PNs responding highly to an odor have elevated spike thresholds and are less-likely to respond at odor offset. At the same time, interconnectivity of PNs with local inhibitory interneurons also contributes to enhancing inverted OFF responses^39^. Our current results suggest that OFF responses in mice are analogous to OFF responses in insect models in both structure and functional consequences. Mitral and tufted cells (MTCs) in mice are analogous to locust PNs and exhibit similar features of intrinsic adaptation^31^. MTCs are also highly interconnected with both local inhibitory granule cells and periglomerular interneurons^40^. Thus, similar cellular and circuit mechanisms may underlie the inversion of odor OFF responses in mouse and locust models. While it is likely that OFF responses in mice are largely caused by a release of odor evoked inhibition, future studies will be required to determine the extent to which this is true.

In contrast to the locust, however, the mouse OB receives extensive feedback from the olfactory cortex, as well as top-down cholinergic and noradrenergic modulation^11–14,41–44^. It has been suggested that persistent odor OFF responses in the mouse are centrally maintained and communicated to the OB via centrifugal feedback^24^. Indeed, differences in circuit mechanisms that maintain OFF responses in mouse and locust may contribute to the notable difference we observed in the persistence of the off response between the two models. Another notable difference between mouse and locust odor responses was the degree of anticorrelation in the OFF. One possible explanation is that increased cortical feedback and top-down control in the mouse olfactory system may actively drive the inverted OFF responses and give rise to more strongly anticorrelated activity compared to the locust model. However, future work is required to test this hypothesis. Nevertheless, the surprising degree of conservation between mouse and locust ON-OFF dynamics, even when observed via very different recording modalities, suggests that analogous circuit mechanisms largely drive and maintain odor OFF responses in both mouse and locust.

Finally, determining the perceptual quality of odor evoked aftereffects and their consequences towards olfactory processing is challenging. While the conservation of persistent and distinct odor aftereffects across species implies that they are a useful feature of olfactory processing, the role of odor aftereffects, their perceptual analogs, and the impact on behavior remain unclear. Suggesting how odor OFF responses may influence behavior, it has been shown that OFF responses in locusts correspond to a behavioral measure of “unsensing”^27^. Additionally, a recent study used behavioral experiments in locust to demonstrate that population odor response trajectories for different odors cluster according to innate and learned valences of odors, indicating that population response structure is related to high-level features of odor perception^45^. While these data suggest that OFF responses are behaviorally meaningful in locusts, no evidence currently exists to suggest how odor OFF responses impact mouse olfactory perception. Future work is therefore required to determine precisely how odor perception in mice and insects is shaped by odor OFF responses, and whether these effects impact odor-guided behavior.

## ACKNOWLEDGMENTS

We thank Dr. Fabrizio Gabbiani for feedback on the manuscript, Pearl Olsen for insect care, and Evelyne Tantry and Elaine Le for mouse care. This research was supported by NSF (1724218, 2021795) and ONR (N00014-19-1-2049, N00014-21-1-2343) grants to B.R., NIH (K99DC019505) to EHM, the McNair Medical institute, NIH awards (NINDS R01NS078294), (NIDDK R01DK109934), and DOD (PR180451-PRMP) awards to B.R.A, and NIH awards (NINDS UF1NS111692) to BRA and JR.

## AUTHOR CONTRIBUTIONS

Conceptualization: BRA and BR; Methodology: DL, EHM, BRA, BR, JR; Formal Analysis: DL, FD, and BR; Investigation: DL, EHM, RK; Data Curation: DL, BR, EHM, CLS, JR; Writing – Original Draft: DL and EHM, Writing – Reviewing and Editing: BRA and BR; Visualization: DL, EHM, and FD; Supervision: BRA, BR and JR; Project Administration: BR; Funding Acquisition: EHM, JR, BR, and BRA

## SUPPLEMENTAL FIGURES AND LEGENDS

**Fig. S1:**
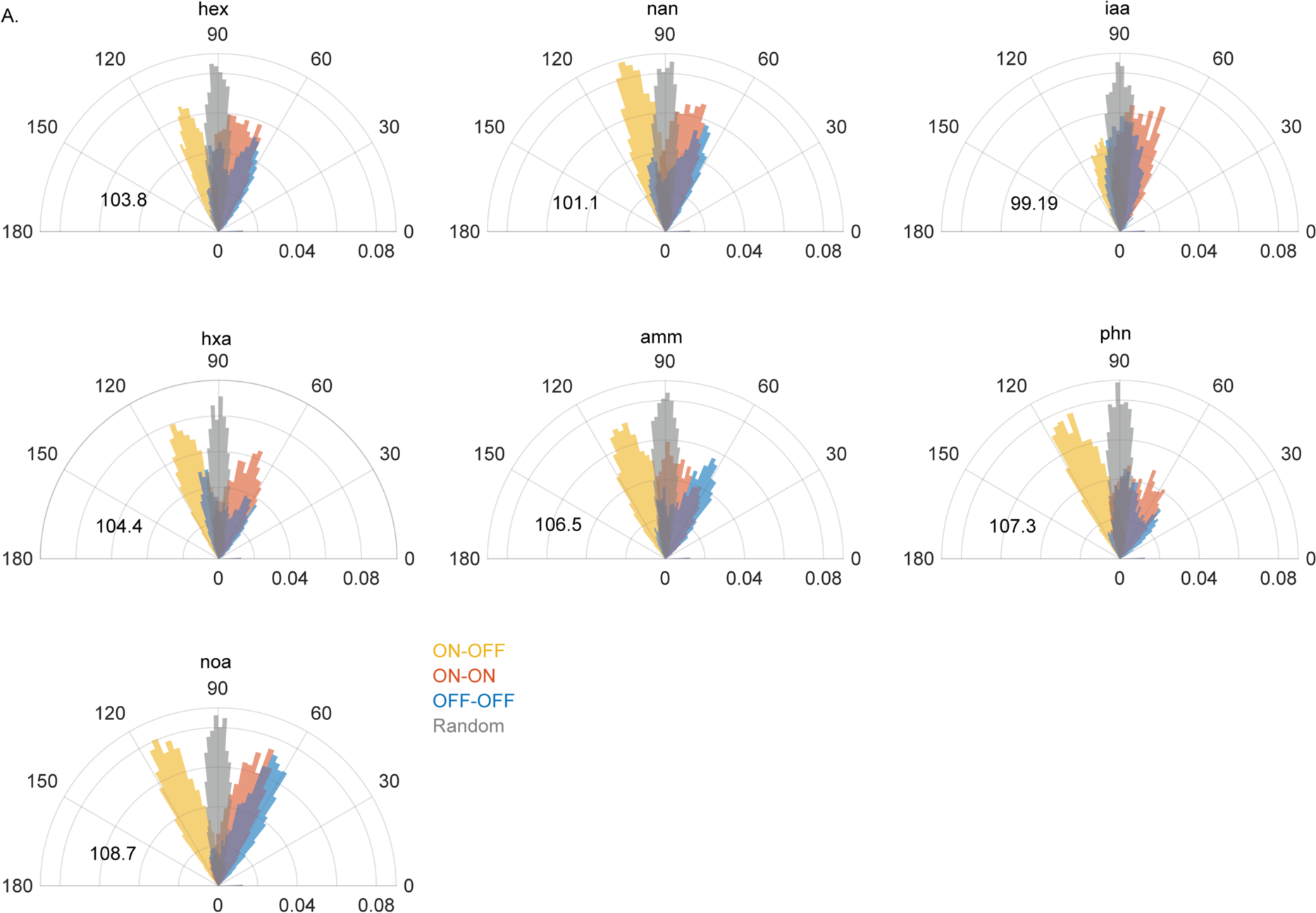
Distribution of angular shifts of PN population responses for remaining odors. Angular shifts in PN population responses comparing each time bin with every other time bin, showing results as probability density (radii) for remaining seven odors not shown in Fig.1. Odors include hexanol (hex), isoamyl acetate (iaa), 1-nonanal (nan), hexanoic acid (hxa), phenethylamine (phn), ammonia (amm), and nonanoic acid (noa). ON period time bins are compared to other ON period time bins (orange) and OFF-period time bins are compared to other OFF period time bins (blue) to show the spread of PN population responses within ON and OFF periods. ON period time bins are compared to OFF period time bins (yellow) to show the angular shift between ON and OFF responses. Points from random vectors (grey) are compared to show the centering of the random distributions around 90°. Median values of ON vs. OFF period angular distributions are shown as black text.

**Fig. S2:**
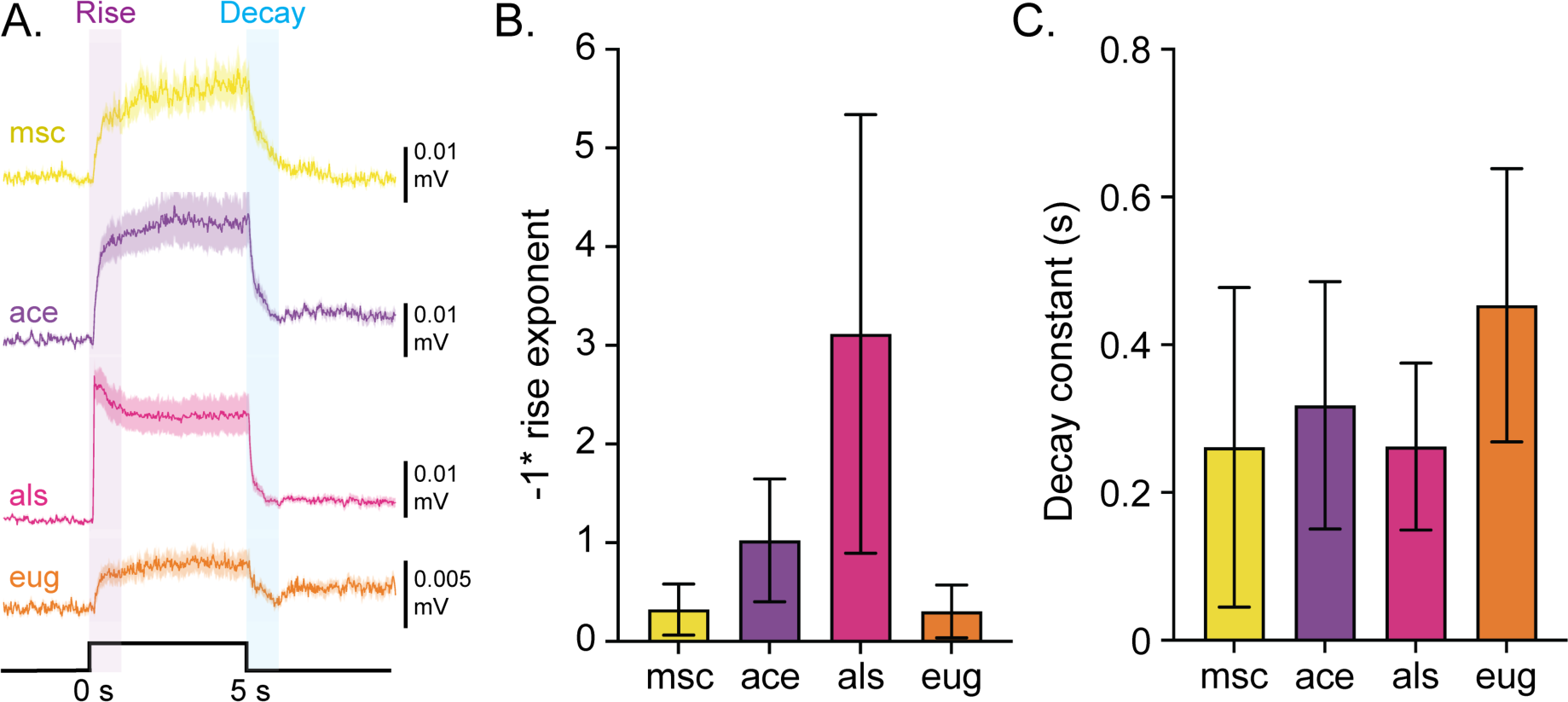
Kinetics of mouse odor presentations. **A.** Example photoionization detector (PID) recordings taken from directly adjacent to the mouse nostril during odor presentations. Thick lines show <u>baseline-subtracted</u> PID traces averaged across 10-20 odor presentations from an example recording session. Shaded areas show SEM. One second time windows after odor onset and offset are highlighted to show data from which rise and decay fits were calculated. **B.** Rise times for each odor across datasets were fit with two-parameter power function f(t) = a * t^b + c. Bars show the mean and SEM of the estimation the exponent b. **C.** Decays were fit with the biexponential function f(t) = a * exp(b * t) + c * exp(d * t). Bars show the mean and SEM of the estimation of the decay constant (-1/b) across imaging sessions.

**Fig. S3:**
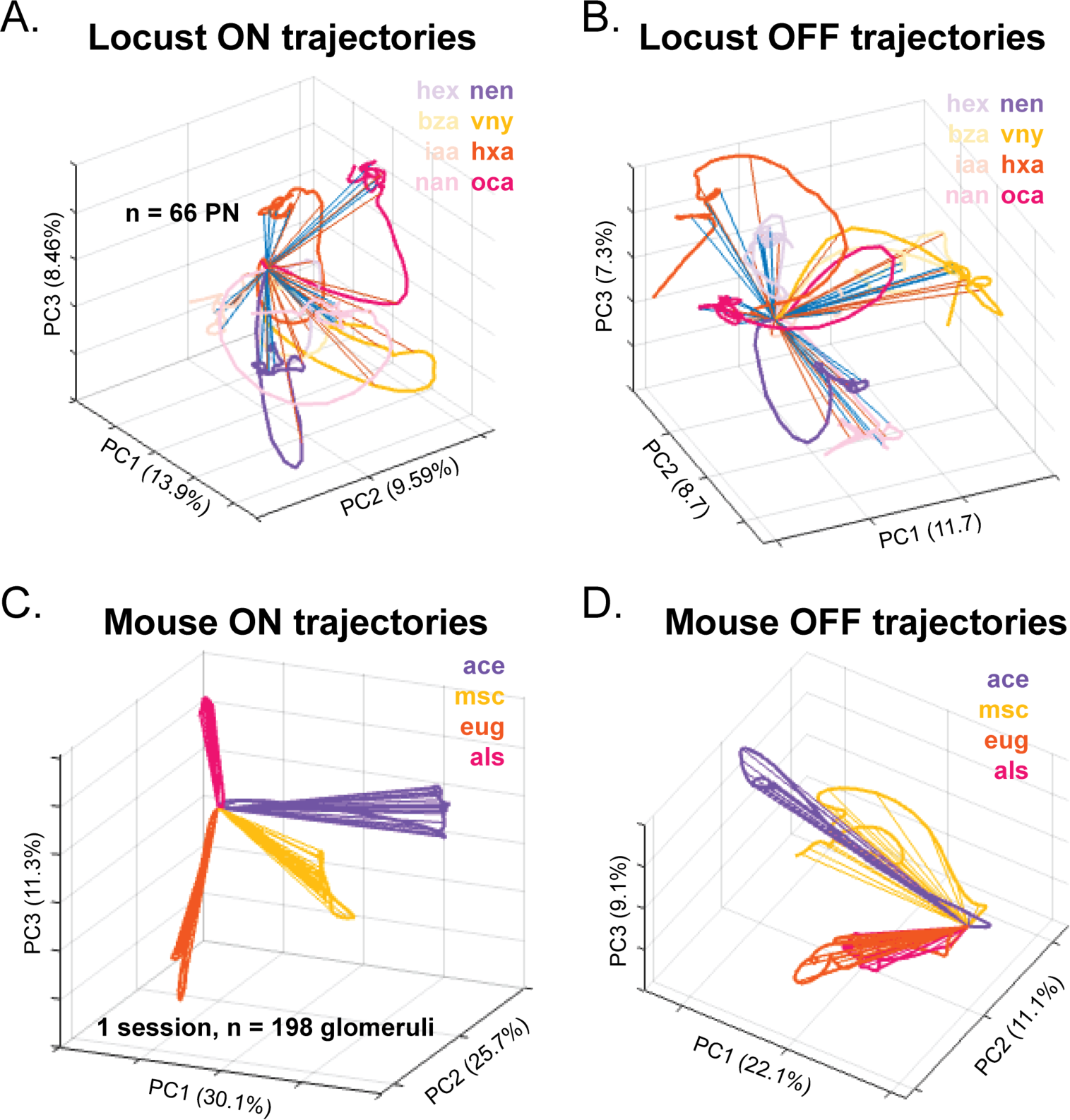
Odor ON and OFF response trajectories projected onto separate principal components for locust and mouse. **A.** Dimensionality-reduced odor-evoked ON responses from the population of locust AL PNs. ON Responses to 8 odors are projected onto the top three principal component axes calculated using only the ON responses. Odors shown include benzaldehyde (bza), hexanoic acid (hxa), hexanol (hex), isoamyl acetate (iaa), 1-nonanal (nan), z-3-nonen-1-ol (nen), octanoic acid (oca), 4-vinyl anisole (vny). **B.** Dimensionality-reduced OFF responses from the population of locust AL PNs. OFF Responses to 8 odors are projected onto the top three principal component axes calculated using only the OFF responses. **C.** Dimensionality-reduced odor-evoked ON responses from a representative imaging session of mouse OB glomeruli. ON Responses to 4 odors are projected onto the top three principal component axes calculated using only the ON responses. Odors presented include methyl salicylate (msc), acetophenone (ace), eugenol (eug), and allyl sulfide (als). **D.** Dimensionality-reduced OFF responses from a representative imaging session of mouse OB glomeruli. OFF responses to 4 odors are projected onto the top three principal component axes calculated using only the OFF responses.

**Fig. S4:**
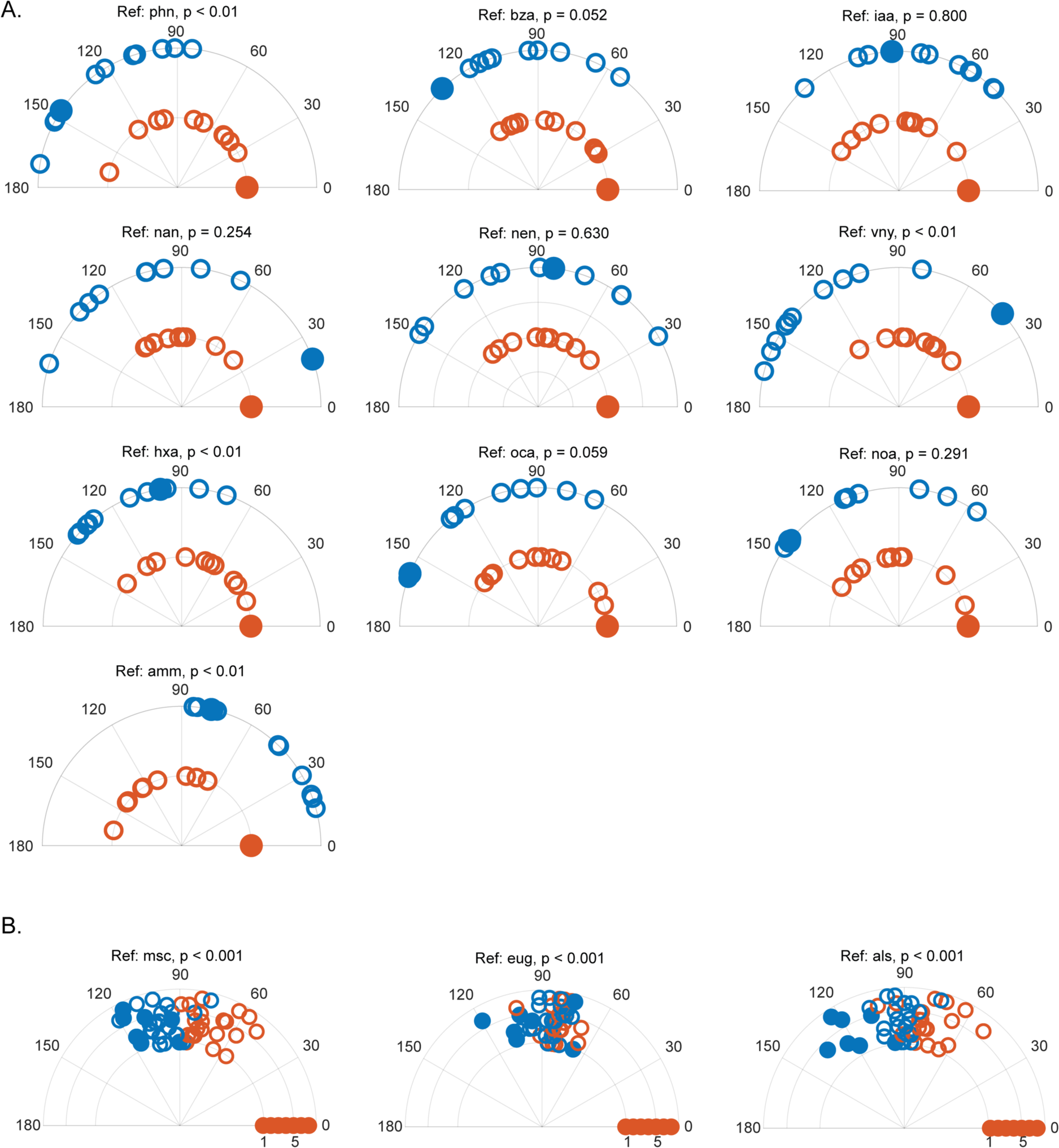
Separation between ON and OFF responses across reference odors. **A.** Polar plots showing the median angle between the locust AL PN response vector for a reference odor (black text) ON response (solid orange circles), ON responses to the 10 other odors (open orange circles), the reference odor OFF response (solid blue circle), and OFF-responses to the 10 other odors (open blue circles). Quantification of angular shifts is shown for 10 reference odors not shown in Figure 3. **B.** Polar plots showing the median angle between the mouse OB glomerular response vector for a reference odor (black text) ON response (solid orange circles), ON responses to the 3 other odors (open orange circles), the reference odor OFF response (solid blue circle), and OFF responses to the 3 other odors (open blue circles). Quantification of angular shifts is shown for 3 reference odors not shown in Figure 3. Radii indicate different imaging sessions (N = 7 sessions from 4 mice).

**Fig. S5:**
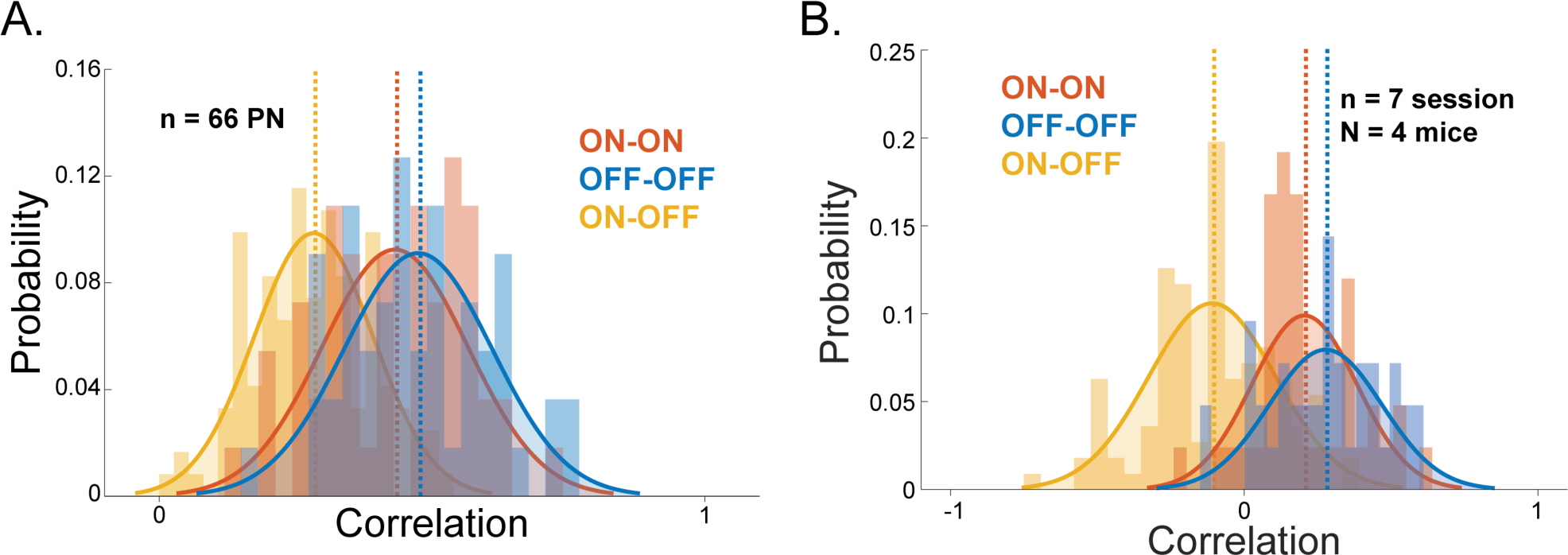
Distributions of correlation values across odor ON and OFF periods in mouse and locust. **A.** From locust data, histograms revealing probability density functions of correlation values comparing ON responses across odors (ON-ON, orange), OFF responses across odors (OFF-OFF, blue), and ON responses across odors to OFF responses across odors (ON-OFF, yellow). Solid line and shaded Gaussian distributions are maximum-likelihood fits of the three correlation distributions. Dashed colored lines show means of each distribution **B.** From mouse data, histograms of correlation values comparing ON responses across odors (ON-ON, orange), OFF responses across odors (OFF-OFF, blue), and ON responses across odors to OFF responses across odors (ON-OFF, yellow). Gaussian distribution and mean (dashed lines) fit using maximum likelihood estimation are also shown.

**Fig. S6:**
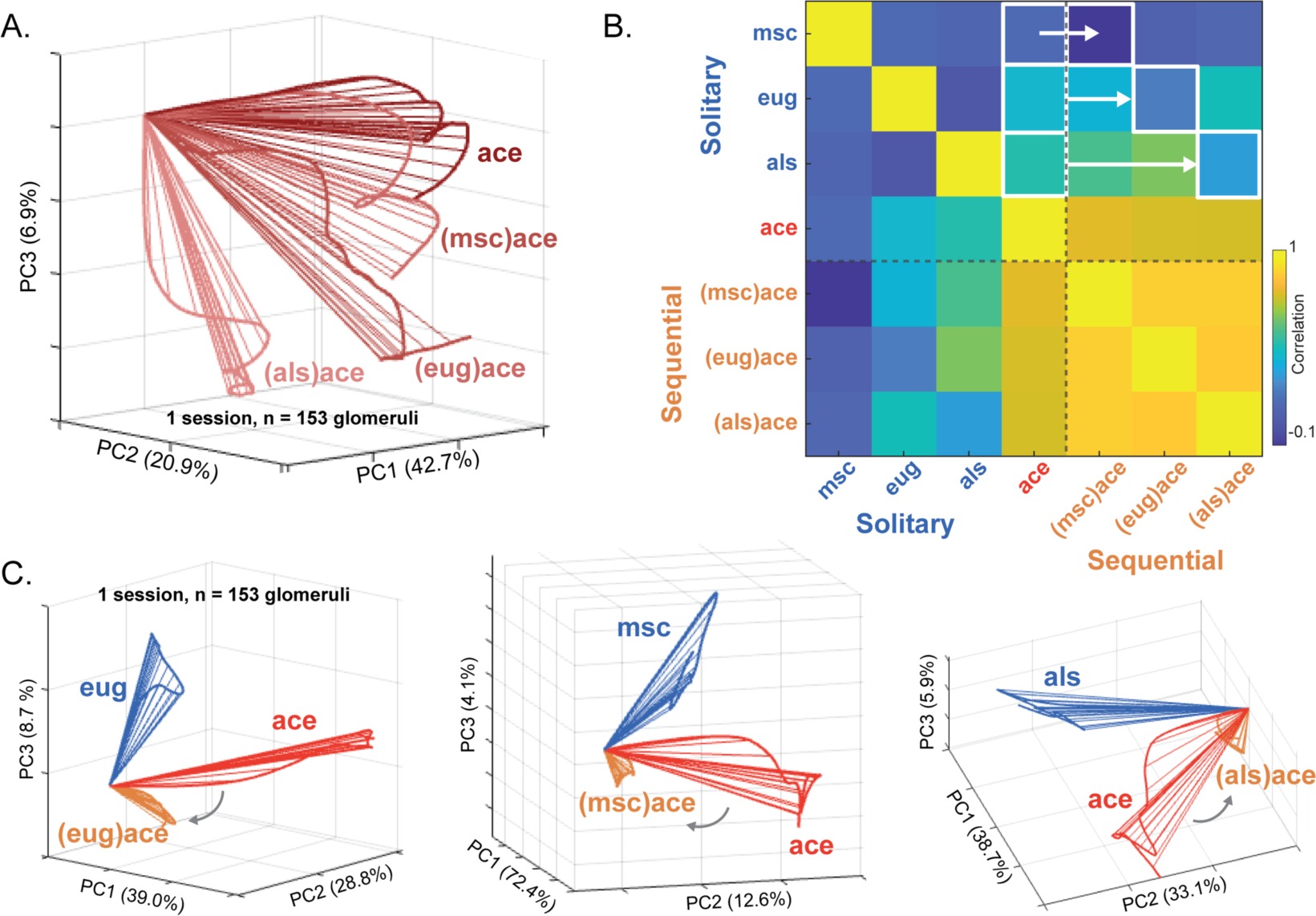
Contrast enhancement in the mouse olfactory bulb generalizes across odors. **A.** Representative dimensionality reduced temporal trajectories showing the response to a solitary presentation of acetophenone (ace, dark red) and responses to ace sequentially presented after methyl salicylate ((msc)ace), eugenol ((eug)ace), and allylsulfide ((als)ace, lighter reds). **B.** Correlation of average glomerular responses during ON periods across solitary presentations of eug, ace, msc, and als as well as sequential presentations of ace following msc ((ace)msc), eug ((eug)ace), and allyl sulfide ((als)ace). Each box represents the correlation of trial averaged glomerular response vectors between odor presentation conditions from a representative imaging session. White boxes and arrows indicate which correlation values from the representative imaging session are used for the quantitative comparison in Figure 6E. To the left of the arrows, boxes highlight correlations between solitary presentations of ace with solitary presentations of other odors (solitary odor vs. solitary ace). To the right of the arrows, boxes highlight correlations between presentations of ace in sequence with other odors and solitary presentations of the other odors (solitary odor vs. sequential (odor)ace). **C.** Representative dimensionality reduced temporal trajectories showing responses to solitary and sequential presentations of ace. (left) Dimensionality reduced ON responses to solitary eug (blue), solitary ace (red), and ace sequentially presented after eug ((eug)ace, orange). (middle) Dimensionality reduced ON responses to solitary msc (blue), solitary ace (red), and ace sequentially presented after msc ((msc)ace, orange). (right) Dimensionality reduced ON responses to solitary als (blue), solitary ace (red), and ace sequentially presented after als ((als)ace, orange).

**Fig. S7:**
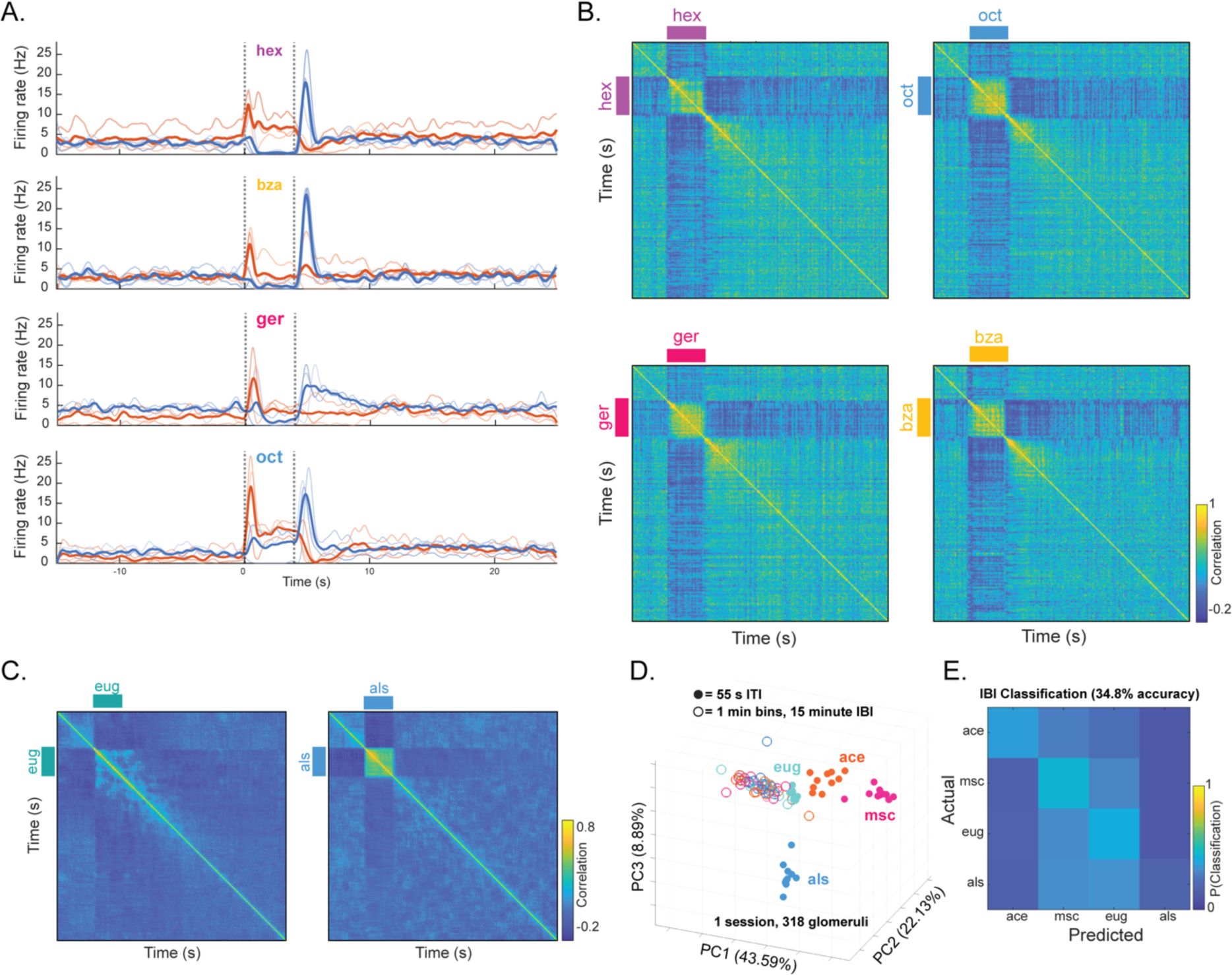
OFF response persistence across odors through intertrial and inter-block intervals. **A.** Trial averaged PSTH traces from representative ON and OFF responsive ONs showing pre-odor, odor-evoked, and post odor activity for odors hexanol (hex), benzaldehyde (bza), geraniol (ger), and octanol (oct) reflecting the remaining odors not shown in Figure 7. **B.** Representative temporal correlations of PN responses to app, cit, ger, and oct, reflecting the remaining odors not shown in Figure 7. Each pixel represents the correlation of the trial averaged PN population response vectors between two 50 ms time bins. **C.** Representative temporal correlations of OB glomerular responses to eug and als. Each pixel represents the correlation of the trial averaged glomerular population response vector at one time point compared to one other time point. **D.** Dimensionality-reduced mouse OB glomerular responses during the 55-second intertrial intervals (ITIs, solid circles) and 15-minute inter-block intervals (IBIs, open circles) plotted on the same three principal component axes. Each point represents 55 seconds of activity between successive trials of the same odor within a block (solid circles) or successive one-minute bins of the intervals between blocks of different odors (open circles). **E.** K-NN classification based on angular distance of the glomerular response vectors during one-minute bins of the 15-minute IBI. The overall classification rate of the glomerular IBI activity was 34.8%.

**Fig. S8:**
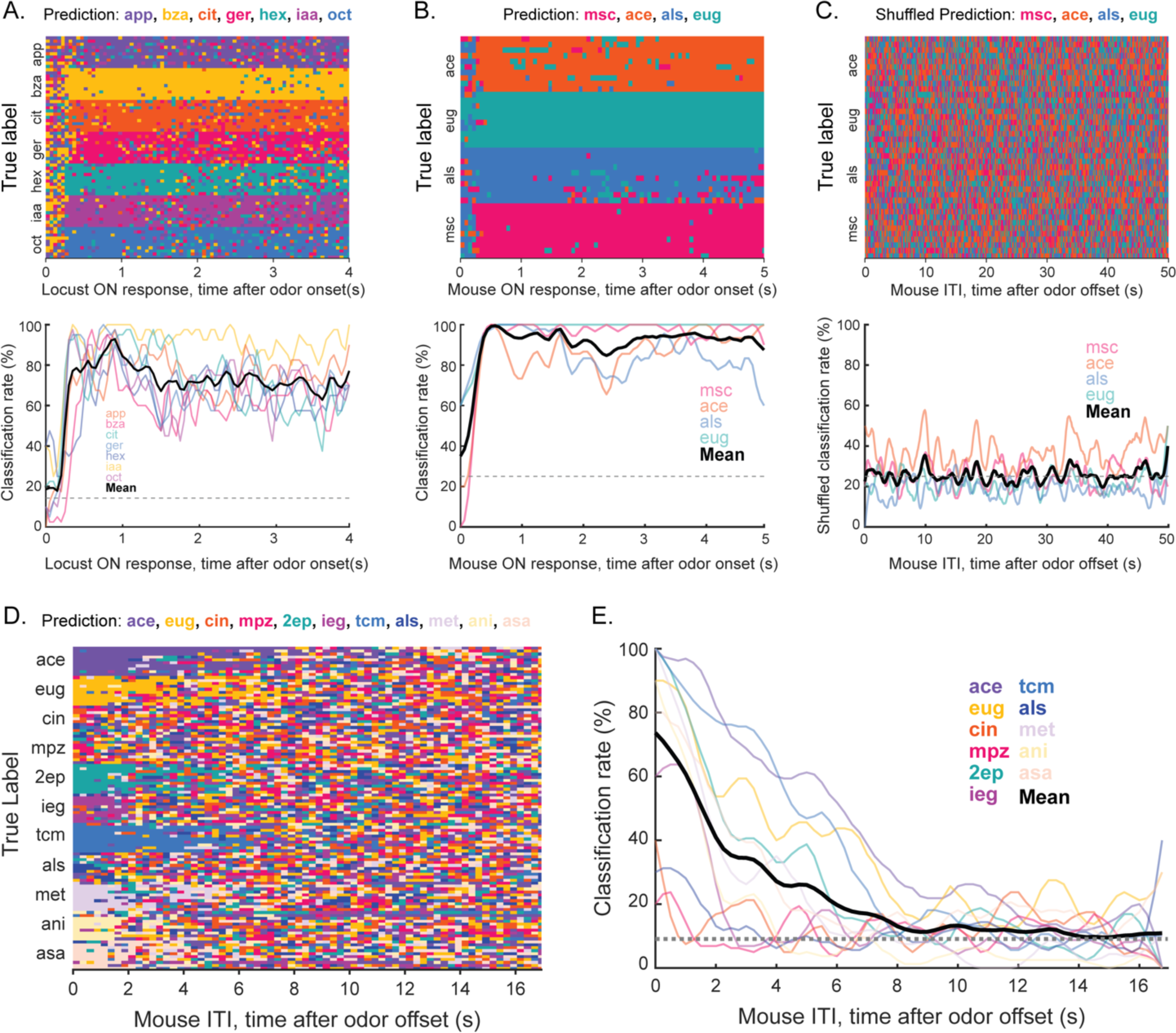
Odor classification over time from locust AL and mouse OB. **A.** (top) Locust data, time-bin-by-time-bin k-NN classifier (k = 10; correlation distance; locusts PN activities) predictions for the odor presentation ON period time bins are shown. Odor predictions at each timepoint are color coded. (bottom) Quantification of classifier accuracy during the odor presentation ON period for each odor (colored transparent lines) and averaged across odors (black line) are shown. The chance-level is identified using the dashed grey line. **B.** (top) Mouse data time-bin-by-time-bin k-NN classifier (*k* = 10; correlation distance; mouse glomerular activities) predictions for the odor presentation ON period time bins are shown. Odor predictions at each timepoint are color coded. (bottom) Quantification of classifier accuracy for the odor presentation ON period (colored transparent lines) and averaged across odors (black line) are shown. The chance level is identified as the dashed grey line. **C.** (top) Results from a statistical control analysis of mouse data where the odor labels in the dataset were randomly shuffled are shown. A time-bin-by-time-bin k-NN classifier (*k* = 10; correlation distance; mouse glomerular activities) was used to generate predictions shown for the post stimulus ITI time bins. Odor predictions at each timepoint are color coded. (bottom) Quantification of classifier accuracy during the ITI time window (colored transparent lines) and averaged across odors (black line) are shown as a function of time. The chance level is identified as the dashed grey line. **D.** Mouse data with eleven odors presented in a random order. Odors included acetophenone (ace), eugenol (eug), cineole (cin), methypyrazine (mpz), 2-ethylpyrizine (2ep), isoeugenol (ieg), transcinnamaldehyde (tcm), allylsufide (als), methylsalycylate (met), anisole (ani), and isoamyl acetate (asa). Time-bin-by-time-bin k-NN classifier (*k* = 10; correlation distance; mouse glomerular activity) predictions of the post stimulus ITI time bins. Odor predictions at each timepoint are color coded. **E.** Quantification of classifier accuracy from D during the ITI time window after random-order odor presentations (colored transparent lines) and averaged across odors (black line). The chance level is identified as the dashed grey line.

**Table S1:** Summary of datasets used in each figure. Locust and mouse datasets used for analyses and figure generation are listed for each figure along with their characteristics.

| Figure | Dataset |
| --- | --- |
| 1 | Locust dataset 1: 66PNs across 15 animals |
| 2c-g | Mouse 2, session 3: 198 glomeruli |
| 2h | 7 mouse datasets: m1, session 1, 73 glomeruli; m2, session1, 210 glomeruli; m2 session 2, 153 glomeruli; m2, session 3, 198 glomeruli; m3, session 1, 286 glomeruli; m4, session 1, 219 glomeruli; m4, session 2, 318 glomeruli |
| 3 | Locust dataset: 66PNs across 15 animals |
| 4 | Mouse 2, session 3: 198 glomeruli |
| 5 | Reanalysis of locust dataset from (cite): 85PNs across 9 animals |
| 6a-d | Mouse 4, session 1: 219 glomeruli |
| 6e, left | 3 Mouse datasets: m3, session 1; m4, session 1; m4, session 2 |
| 6e, right | 4 Mouse datasets: m1, session 1; m2, session 1; m2, session 2; m2, session 3 |
| 7a, c, e, f, i | Reanalysis of locust dataset from<sup>27</sup>: 85PNs across 9 animals |
| 7b,d,g,h,j,k | Mouse 4, session 2: 318 glomeruli |
| S1 | Locust dataset 1: 66PNs across 15 animals |
| S2 | 4 Mouse datasets: m1, session 1; m2, session 1; m2, session 2; m2, session 3 |
| S3a, b | Locust dataset 1: 66PNs across 15 animals |
| S3c, d | Mouse 2, session 3: 198 glomeruli |
| S4a | Locust dataset #1: 66PNs across 15 animals |
| S4b | 7 mouse datasets: m1, session 1; m2, session 1; m2, session 2; m2, session 3; m3, session 1; m4, session 1; m4, session 2 |
| S5a | Locust dataset #2: 85PNs across 9 animals |
| S5b | Mouse 2, session 3: 198 glomeruli |
| S6 | Mouse 2, session 2: 153 glomeruli |
| S7a-b | Reanalysis of locust dataset from<sup>27</sup>: 85PNs across 9 animals |
| S7c-d | Mouse 4, session 2: 318 glomeruli |
| S8a | Reanalysis of locust dataset from<sup>27</sup>: 85PNs across 9 animals |
| S8b-c | Mouse 4, session 2: 318 glomeruli |
| S8d-e | Mouse 8, session 1: 175 glomeruli |

## METHODS

### Resource Availability

#### Lead contact

- Further information and requests for resources and reagents should be directed to and will be fulfilled by the lead contacts, Ben Arenkiel, and Barani Raman.

#### Data and code availability

- Locust electrophysiology and mouse calcium imaging data will be deposited in Figshare and made publicly available as of the date of publication. Accession numbers will be listed in the key resources table.
- Original code used to analyze data and generate figures in this paper will be deposited in Figshare and become publicly available as of the date of publication. DOIs will be listed in the key resources table.
- Any additional information required to reanalyze the data reported in this paper is available from the lead contact upon request.

### Experimental Model and Study Participant Details

#### Mice

All experimental procedures were approved by the Baylor College of Medicine Institutional Animal Care and Use Committee. Thy1-GCamp6f (Jax laboratories #025393) mice, age two months to four months, of both sexes, were used for two-photon imaging experiments. Mice were housed in a standard 12-hour light/dark cycle and had *ad libitum* access to food and water.

#### Locusts

Post-fifth instar adult locusts (*Schistocerca americana*) were reared in a crowded colony with a 12-hour light-dark cycle. Both male and female were used for electrophysiological experiments.

### Method Details

#### Cranial window surgery

Chronic cranial windows were created in mice by removing a 4 mm diameter section of skull over the OB and inserting a glass coverslip. Before surgery mice were treated with 5 mg/kg Meloxicam. Anesthesia was induced and maintained with isoflurane during the surgical procedure. After induction of anesthesia, the scalp was injected subcutaneously with 0.05 mL bupivacaine, then cleaned and removed over the OB and dorsal skull. A 0.0.16’’ thick stainless-steel shim (McMaster-Carr, A370-974) was centered over the OB and attached to the exposed skull with dental cement (C&B Metabond). A 4 mm diameter piece of skull was removed by carefully drilling though the skull. A 4 mm glass coverslip (Warner instruments) was placed over the exposed brain and sealed in place with tissue adhesive (3M Vetbond). The sealed window was then stabilized with high-viscosity cyanoacrylate superglue (Loctite). After the glue fully cured, the coverslip was protected with a cap of Kwik-Cast silicone elastomer (World Precision Instruments) which was removed immediately before imaging.

#### Odor delivery to mice

All imaging was performed on head-fixed, awake mice on a running wheel. Mice were head fixed using a custom headplate^47^ designed to attach to the stainless-steel shim implanted on the skull during the cranial window surgery. Mice were habituated to the head fixation and imaging setup for at least 30 minutes prior to testing each day. For odor delivery, a multi-channel olfactometer^48^ was placed 6 cm in front of the mouse. The olfactometer provided a constant stream of room air into which experimental odors were injected. Odors were mixed into the central airstream before delivery to the mouse by an eductor positioned at the output of the olfactometer. Odors for mouse imaging experiments were obtained from Sigma and included methyl salicylate (msc), allyl sulfide (als), acetophenone (ace), and eugenol (eug), diluted to 10% by volume in mineral oil. Odors were further diluted to ∼1% of their initial concentration by injection into the central airstream (∼8 L/min flow rate). Odor delivery was controlled and synchronized with imaging via custom LabVIEW software. The timing of odor onset and offset was verified with PID recordings (200B miniPID, Aurora Scientific) made adjacent to the mouse nostril **(Fig. S2).** For solitary odor presentation experiments, odors were delivered individually for 5 seconds via injection into the central air stream. Each odor was repeated 10 times in a block with a 55 seconds intertrial interval between presentations within a block and 15 minutes between blocks of different odors. For sequential presentation blocks, odors were presented in pairs such that the first odor was presented for 5 seconds followed by a 1 second gap and then 5 seconds of the second odor. Odor pairs were presented in blocks of 10 with 49 seconds between trials within a block and 15 minutes between blocks of different odor pairs. For random order odor presentations (**Fig. S8 D, E**), odors were presented one second with a 19 second intertrial interval. 11 different odors were presented in a random order over 480 trials with no block structure (i.e., without 15 minutes of no-odor presentation). For experiments using 11 odors, odors (Sigma) included acetophenone (ace), eugenol (eug), cineole (cin), methypyrazine (mpz), 2-ethylpyrizine (2ep), isoeugenol (ieg), transcinnamaldehyde (tcm), allylsufide (als), methylsalycylate (met), anisole (ani), and isoamyl acetate (asa), all diluted to 1:10 by volume and further diluted ∼1:100 during delivery. Odors were scavenged after delivery by a constant vacuum positioned 10 cm behind the mouse in line with the air stream of the olfactometer.

#### Two-photon imaging

Two-photon imaging was performed on a ThorLabs/Janelia 2P-RAM mesoscope^28^. The laser wavelength was set to 920 nm to image GCaMP6f signals. Imaging parameters were controlled with ScanImage software. To maximize frame rates in each imaging session, fields of view were defined for acquisition that included a single plane visualizing only the dorsal surface of bilateral OBs (1800 um long x ∼600um –wide fields of view that were tiled to cover a total field of view that was 1800-2500 um wide). Images were acquired continuously throughout experiments (during odor presentations, intertrial, and inter-block intervals), with 5um/pixel resolution at the fastest possible frame rate allowed by the imaging parameters (15-18 Hz). After imaging, videos were motion and raster-corrected and glomerular ROIs were manually defined with custom software. Following definition of glomerular ROIs, fluorescence traces were extracted from ROIs, imported to MATLAB, converted to dF/F, and median filtered to remove any persistent motion artifacts.

#### Locust electrophysiology

Locusts were immobilized with both antennae intact. Then the primary olfactory region of their brain, the antennal lobe (AL), was exposed, desheathed, and perfused with room temperature saline. Extracellular multiunit recordings of projection neurons (PNs) were performed with a 16-channel, 4×4 silicon probe (NeuroNexus) that was superficially inserted in the AL. Prior to each experiment, all probes were electroplated with gold to achieve impedances in the range of 200 to 300 kΩ. The recordings were acquired with a custom 16-channel amplifier (Biology Electronics Ship; Caltech, Pasadena, CA). The signals were amplified with a 10k gain, bandpass filtered (0.3 to 6 kHz), and sampled at 15 kHz using a LabView data acquisition system. A visual demonstration of this protocol is available online^49^.

#### Odor delivery to locusts

Odor stimuli were delivered using a standard protocol previously described in our earlier work^45^. The following odor panel was used for electrophysiological experiments: ammonia (amm), apple (app), benzaldehyde (bza), citral (cit), geraniol (ger), hexanol (hex), hexanoic acid (hxa), isoamyl acetate (iaa), 1-nonanal (nan), z-3-nonen-1-ol (nen), nonanoic acid (noa), octanoic acid (oca), octanol (oct), phenethylamine (phn), and 4-vinyl anisole (vny). All odors were diluted in mineral oil to 1% v/v concentration and sealed in 60mL glass bottles with an air inlet and outlet. A pneumatic picopump (WPI Inc., PV-820) was used to deliver 0.1L/min of air to the odor bottle. The injected air displaced air containing diluted odorant in the bottle headspace, which was subsequently mixed with a desiccated 0.75L/min carrier air stream directed towards the locust antennae. A vacuum funnel placed behind the locust preparation continuously removed delivered odors.

### Quantification and Statistical Analysis

#### Summary of locust and mice datasets

Two locust datasets were used to generate **Figs. 1, 3, 5**, and **7**. The dataset used for generating **Figs. 5** and **7** were part of an earlier published dataset^52^. Seven mouse datasets were used to generate **Fig. 2**, **4**, **6**, and **7**. All datasets are summarized in **Supplemental Table 1**.

#### PN spike sorting

To obtain single-unit PN responses, spike sorted was performed offline using four recording channels and conservative statistical principles^50^. Spikes belong to single PNs were identified as described in earlier work^51^. The following criteria were used to identify single units: cluster separation > 5 x noise standard deviations, total number of spikes within 20 ms inter-spike interval < 6.5% of total spikes, and spike waveform variance < 6.5 x noise standard deviations. In total, 66 PNs from 15 locusts were identified for data shown in **Figs. 1** and **3**; and 85PNs were identified from 9 locusts for data shown in **Figs**. **5** and **7**.

#### Time-bin-by-time-bin correlation analysis

Each pixel or matrix element in time-bin-by-time-bin correlation plots (**Figs. 1E**, **2F**, **7C**, **D**) indicates the correlation value between neural activity vectors observed in the *i*^th^ and *j*^th^ time bins. All time-bin-by-time-bin correlation analyses were computed using high-dimensional response vectors. Correlations were calculated as:

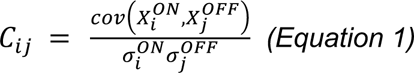

Here, *X* is a n-dimensional activity vector, *i* and *j* represent time bins, 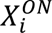 represents the population activity vector in the *i^th^*time bin during the ON period, 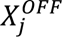 represents the population activity vector in the *j^th^*time bin during the OFF period, 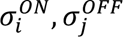 are standard deviations of spiking activities during ON and OFF periods during the i^th^ and j^th^ time bins respectively.

For locust data, PN spikes were binned in 50 ms non-overlapping time bins, and spike counts of different PNs were concatenated to obtain a n-dimensional population spike count vector (where n = 66 PNs for **Fig. 1E**; n = 85 PNs for **Fig. 7C**). For mouse data, dF/F values were taken to be a n-dimensional population glomerular activity vectors (where n=198 glomeruli for **Fig. 2F**; n = 318 glomeruli for **Fig. 7D**).

#### Time averaged correlations analysis

All time averaged correlation analyses were computed using high-dimensional response vectors. Correlations were calculated as:

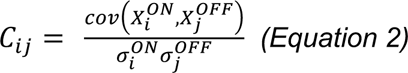

Here, *X^ON^*, *X^OFF^* are trial- and time-averaged high-dimensional activity vectors for the ON and OFF periods, respectively, respectively.

For locust data, PN spikes during the 4 s odor stimulation were summed and averaged across ten trials to obtain a n-dimensional population PN average spike count vector (where: n = 66 PNs **for Fig. 3C**; n = 85 PNs for **Fig. 5D**). For mouse data, dF/F values were averaged across the 10 trials, and then averaged across the frames for the 5 second odor stimulation duration to obtain a n-dimensional population glomerular activity vector (where n = 198 glomeruli for **Fig. 4C**; n = 219 glomeruli for **Fig. 6D**).

#### Visualization of response trajectories through PCA

To visualize high dimensional AL PN spike counts and OB glomerular dF/F signals, we used a linear principal component analysis. For locust data, the spike counts for each PN in 50 ms time bin during the 4 second stimulation window were averaged across trials and concatenated across PNs to generate a *n*-dimensional vector (where *n* = 66 PNs for **Fig. 1D**; trial-averaged neural trajectories). The high-dimensional PN spike count vector was projected onto the top three eigenvectors of the data covariance matrix.

For mouse data, dF/F values at each imaging frame (captured at 15-19 Hz) during the 5 second stimulation window were averaged across trials and concatenated across glomeruli to generate a n-dimensional vector (where n = 198 glomeruli for **Fig. 2E**). The high-dimensional glomerular response vector was projected onto the top three eigenvectors of the data covariance matrix.

#### Visualization of ITI and IBI neural activity vectors through PCA

To visualize high dimensional AL PN spike counts and OB glomerular dF/F signals during intertrial intervals (ITI) and inter-block intervals (IBI) we used a linear principal component analysis. For locust data, the ITI was defined as the 21 seconds immediately following the odor offset. The spike counts for each PN were averaged across the 20 second ITI period. The resulting n-dimensional PN spike count vector (where n = 85 PNs for **Fig. 7E**) was computed for each trial and visualized after PCA dimensionality reduction. The data points were colored based on the odor that preceded the ITI analysis window. For mouse data, the ITI was defined as the 50 seconds immediately following the odor offset. The dF/F values were averaged across the 50 second ITI period for each glomerulus and the resulting n-dimensional glomerular activity vector (where n = 318 glomeruli for **Fig. 7G**) represented the glomerular activity following the odor offset for a given trial. Glomerular activity following odor termination in different trials were visualized after PCA dimensionality reduction, and the data points were colored based on the odor that preceded the ITI period.

For mouse data, the IBI was defined as the 15-minute non-odor stimulation period between blocks of odor stimulation. The dF/F values were binned into 1-minute non-overlapping time bins across the 15-minute IBI period and concatenated. The resulting n-dimensional glomerular activity vector (where n = 318 glomeruli for **Fig. S7D**) represented the glomerular activity during the entire IBI period. The IBI activity vectors were visualized after dimensionality reduction and the data points colored based on the odor that was presented in the block of trials that preceded the IBI window.

#### Distribution of angles between high-dimensional response vectors

Vector angles were calculated between trial-averaged PN activity for each 50 ms time bin in the 4 second ON period (**Fig. 1F**, **2G**). The resulting vector angles were subsequently binned according to the ON-ON, ON-OFF, or OFF-OFF period depending on the time window from which the PN vectors were picked. A similar analysis was carried out for mouse OB glomerular responses using dF/F values from each imaging frame. As a control, the vector angles were calculated from high-dimensional normally distributed random vectors. For hierarchical clustering, we first calculated the summed spike counts during 4 second odor presentations for each individual PN and generated a single vector (*n* = 66 PNs) for each odor. Similarly, dF/F values for each glomerular ROI in the mouse OB imaging datasets were averaged across the duration of an odor ON or OFF period. Dendrograms were generated using the angular distance between respective average odor responses. The cluster tree was created in such a way that the furthest pairwise distance between any two samples assigned to an individual cluster was minimized.

#### Classification analysis of average ITI periods

To classify ITI periods, we used a k-Nearest Neighbor (k-NN) classification algorithm. For locust data, the ITI was defined as the 20 seconds immediately following the odor offset. The spike counts for each PN were binned in 50 ms non-overlapping time bins and averaged across the 20 second ITI period. The resulting n-dimensional PN spike count vector (where n = 85 PNs for **Fig. 7F**) represented the PN activity following the odor offset for a given trial. Then the PN spike count vectors were concatenated across the 10 trials and 7 odors, resulting in 70 trials/vectors for classification. The angular distance of the resulting high-dimensional PN spike count matrix was used as a distance measure and the most represented odorant amongst the 10 nearest neighbors was the label assigned to each trial (i.e., 10-nearest neighbor approach; leave-one-trial-out-validation).

For mouse data, the ITI was defined as the 50 seconds immediately following the odor offset. The dF/F values were averaged across the 50 second ITI period for each glomerulus and the resulting n-dimensional glomerular activity vector (where n = 318 glomeruli for **Fig. 7H**) represented the glomerular activity following the odor offset for a given trial. Then the glomerular activity vectors were concatenated across the 10 trials and 4 odors, resulting in 40 trials/vectors for classification (leave-one-trial-out validation). The angular distance of the resulting high-dimensional glomerular activity matrix was used to classify each trial based on the most represented odorant amongst the 50 nearest neighbors (i.e., 50 nearest neighbors). A larger *k*-value was used compared to the *k*-value used for locust ITI classification to account for the slower calcium indicator dynamics.

#### Classification analysis of average IBI periods

To classify IBI periods, we used a similar k-Nearest Neighbor (k-NN) classification algorithm. For mouse data, the IBI was defined as the 15-minute non-odor stimulation period between blocks of odor stimulation. The dF/F values were averaged over 1-minute non-overlapping time bins across the 15-minute IBI period, resulting in n-dimensional glomerular activity vector (where n = 318 glomeruli for **Fig. S7D**). Then the glomerular activity vectors were concatenated across the 4 odors, resulting in 60 trials for classification (leave-one-trial-out cross-validation; *50-nearest neighbor approach*).

#### Time bin by time bin ITI classification analysis

For locust data, the ITI was defined as the 21 seconds immediately following the odor offset. The spike counts for each PN were binned in 200 ms non-overlapping time bins. The resulting n-dimensional PN spike count vector (where n = 85 PNs for **Fig. 7I**) represented antennal lobe ensemble activity following the odor offset. Then the PN spike count vectors were concatenated across the 7 odors and 10 trials, resulting in 7350 time bins/vectors for classification. The odor label for each time bin was determined using a k-nearest neighbor classifier (k = 10) with leave-one-trial-out validation. A correlation distance between high-dimensional vectors was used as the metric to determine the nearest neighbors. To be rigorous and conservative, for a given trial, the 10-nearest neighbors from only other trials were used to classify (leave-one-trial-out validation). For repeated mouse data, the ITI was defined as the 50 seconds immediately following the odor offset. The dF/F values were averaged across the 10 trials for each glomerulus were averaged every 4 frames in a non-overlapping fashion and the resulting n-dimensional glomerular activity vector (where n = 318 glomeruli for **Fig. 7J**) represented the glomerular activity following the odor offset. Then the glomerular activity vectors were concatenated across the 4 odors and 10 trials, resulting in approximately 8,000 time-bins/vectors for classification. Again, the odor label for each time bin was determined using a k-nearest neighbor classifier (k = 10) with leave-one-trial-out validation. A correlation distance between high-dimensional glomerular vectors was used as the metric to determine the nearest neighbors. To be rigorous and conservative, for a given trial, the 10-nearest neighbors only from other trials were used to classify (leave-one-trial-out validation).

For the analysis of OB dataset that included random presentation of odorants (**Fig. S8D, E**), the ITI was defined as the 17 seconds immediately following the odor offset. The dF/F values for each glomerulus were averaged every 4 frames in a non-overlapping fashion and the resulting n-dimensional glomerular activity vector (where n = 286 glomeruli for **Fig. S8D** and **E**) represented the temporal activity following the odor offset. Then the glomerular activity vectors were concatenated across the 11 odors and first 10 trials for each odor, resulting in 7480 time-bins/vectors for classification. Ten nearest neighbor classification analysis with leave-one-trial-out validation was performed to assign class labels to each glomerular activity vector.

#### Shuffled statistical control analysis

A statistical control analysis, where odor labels were randomly shuffled, was performed on mouse datasets where each odorant was repeatedly presented for ten trials. The ITI was defined as the 50 seconds immediately following the odor offset. The dF/F values for each glomerulus representing the temporal activity following the odor offset were used as the n-dimensional glomerular activity vector (where n = 318 glomeruli for Fig. **S8C**), but the odor label for each time bin/vector was pseudo-randomly generated. Then the glomerular activity vectors were concatenated across the 4 odors and 10 trials, resulting in 16,000 time-bins/vector for classification. Ten nearest neighbor classification analysis with leave-one-trial-out validation was performed to assign class/odor labels to each glomerular activity vector.

#### Time bin by time bin IBI classification analysis

To classify IBI periods, we used a k-Nearest Neighbor (k-NN) classification algorithm. For mouse data, the IBI was defined as the 15-minute non-odor stimulation period between blocks of odor stimulation. The dF/F values per imaging frame during the 15-minute IBI period were calculated and the resulting n-dimensional glomerular activity vector (where n = 318 glomeruli for **Fig. 7K**) represented the glomerular activity during the entire IBI period. Then the glomerular activity vectors were concatenated across the 4 odors, resulting in 72,000 time bins/vectors for classification. The angular distance of the resulting high-dimensional glomerular activity matrix was used to classify each vector using the most common odor label of the nearest *k* points (where *k* = 50).

### Resources Table

| REAGENT or RESOURCE | SOURCE | IDENTIFIER |
| --- | --- | --- |
| **Deposited data** |  |  |
| Locust electrophysiology data | This study | DOI: |
| Mouse calcium imaging data | This study | DOI: |
| **Experimental models: Organisms/strains** |  |  |
| Mouse/ Thy1-GCaMP6f (5.11) | Jax laboratories | Strain #: 025393 |
| **Software and algorithms** |  |  |
| MATLAB | MathWorks | N/A |

## REFERENCES

1. Hildebrand, J. G. & Shepherd, G. M. Mechanisms of olfactory discrimination: converging evidence for common principles across phyla. Annu. Rev. Neurosci. 20, 595–631 (1997).

2. Ache, B. W. & Young, J. M. Olfaction: diverse species, conserved principles. Neuron 48, 417–430 (2005).

3. Buck, L. & Axel, R. A novel multigene family may encode odorant receptors: a molecular basis for odor recognition. Cell 65, 175–187 (1991).

4. Sato, K. et al. Insect olfactory receptors are heteromeric ligand-gated ion channels. Nature 452, 1002–1006 (2008).

5. Spors, H. & Grinvald, A. Spatio-temporal dynamics of odor representations in the mammalian olfactory bulb. Neuron 34, 301–315 (2002).

6. Friedrich, R. W. & Laurent, G. Dynamic optimization of odor representations by slow temporal patterning of mitral cell activity. Science 291, 889–894 (2001).

7. Szyszka, P., Gerkin, R. C., Galizia, C. G. & Smith, B. H. High-speed odor transduction and pulse tracking by insect olfactory receptor neurons. Proc. Natl. Acad. Sci. 111, 16925–16930 (2014).

8. Masse, N. Y., Turner, G. C. & Jefferis, G. S. Olfactory information processing in Drosophila. Curr. Biol. 19, R700–R713 (2009).

9. Jeanne, J. M. & Wilson, R. I. Convergence, divergence, and reconvergence in a feedforward network improves neural speed and accuracy. Neuron 88, 1014–1026 (2015).

10. Olsen, S. R., Bhandawat, V. & Wilson, R. I. Divisive normalization in olfactory population codes. Neuron 66, 287–299 (2010).

11. Zaborszky, L., Carlsen, J., Brashear, H. R. & Heimer, L. Cholinergic and GABAergic afferents to the olfactory bulb in the rat with special emphasis on the projection neurons in the nucleus of the horizontal limb of the diagonal band. J. Comp. Neurol. 243, 488–509 (1986).

12. Cansler, H. L., Denson, H. B. & Wesson, D. W. Centrifugal innervation of the olfactory bulb: a reappraisal. eNeuro 6, (2019).

13. Markopoulos, F., Rokni, D., Gire, D. H. & Murthy, V. N. Functional Properties of Cortical Feedback Projections to the Olfactory Bulb. Neuron 76, 1175–1188 (2012).

14. Boyd, A. M., Sturgill, J. F., Poo, C. & Isaacson, J. S. Cortical Feedback Control of Olfactory Bulb Circuits. Neuron 76, 1161–1174 (2012).

15. Linster, C., Henry, L., Kadohisa, M. & Wilson, D. A. Synaptic adaptation and odor-background segmentation. Neurobiol. Learn. Mem. 87, 352–360 (2007).

16. Kay, L. M. & Laurent, G. Odor-and context-dependent modulation of mitral cell activity in behaving rats. Nat. Neurosci. 2, 1003–1009 (1999).

17. Suh, E., Bohbot, J. D. & Zwiebel, L. J. Peripheral olfactory signaling in insects. Curr. Opin. Insect Sci. 6, 86–92 (2014).

18. Imai, T., Sakano, H. & Vosshall, L. B. Topographic mapping—the olfactory system. Cold Spring Harb. Perspect. Biol. 2, a001776 (2010).

19. Martelli, C., Carlson, J. R. & Emonet, T. Intensity invariant dynamics and odor-specific latencies in olfactory receptor neuron response. J. Neurosci. 33, 6285–6297 (2013).

20. Arevian, A. C., Kapoor, V. & Urban, N. N. Activity-dependent gating of lateral inhibition in the mouse olfactory bulb. Nat. Neurosci. 11, 80 (2008).

21. Sahay, A., Wilson, D. A. & Hen, R. Pattern separation: a common function for new neurons in hippocampus and olfactory bulb. Neuron 70, 582–588 (2011).

22. Gschwend, O. et al. Neuronal pattern separation in the olfactory bulb improves odor discrimination learning. Nat. Neurosci. 18, 1474–1482 (2015).

23. Thompson, P. & Burr, D. Visual aftereffects. Curr. Biol. 19, R11–R14 (2009).

24. Patterson, M. A., Lagier, S. & Carleton, A. Odor representations in the olfactory bulb evolve after the first breath and persist as an odor afterimage. Proc. Natl. Acad. Sci. 110, E3340–E3349 (2013).

25. Barlow, H. A theory about the functional role and synaptic mechanism of visual after-effects. Vis. Coding Effic. 363375, (1990).

26. Kohn, A. Visual adaptation: physiology, mechanisms, and functional benefits. J. Neurophysiol. 97, 3155–3164 (2007).

27. Saha, D. et al. Engaging and disengaging recurrent inhibition coincides with sensing and unsensing of a sensory stimulus. Nat. Commun. 8, 1–19 (2017).

28. Sofroniew, N. J., Flickinger, D., King, J. & Svoboda, K. A large field of view two-photon mesoscope with subcellular resolution for in vivo imaging. Elife 5, e14472 (2016).

29. Rinberg, D., Koulakov, A. & Gelperin, A. Speed-accuracy tradeoff in olfaction. Neuron 51, 351–358 (2006).

30. Virsu, V. & Laurinen, P. Long-lasting afterimages caused by neural adaptation. Vision Res. 17, 853–860 (1977).

31. Vinograd, A., Livneh, Y. & Mizrahi, A. History-dependent odor processing in the mouse olfactory bulb. J. Neurosci. 37, 12018–12030 (2017).

32. Whitmire, C. J. & Stanley, G. B. Rapid sensory adaptation redux: a circuit perspective. Neuron 92, 298–315 (2016).

33. Huk, A. C., Ress, D. & Heeger, D. J. Neuronal Basis of the Motion Aftereffect Reconsidered. Neuron 32, 161–172 (2001).

34. Mather, G., Pavan, A., Campana, G. & Casco, C. The motion aftereffect reloaded. Trends Cogn. Sci. 12, 481–487 (2008).

35. Anstis, S., Verstraten, F. A. & Mather, G. The motion aftereffect. Trends Cogn. Sci. 2, 111–117 (1998).

36. Szyszka, P., Ditzen, M., Galkin, A., Galizia, C. G. & Menzel, R. Sparsening and temporal sharpening of olfactory representations in the honeybee mushroom bodies. J. Neurophysiol. 94, 3303–3313 (2005).

37. Chalasani, S. H. et al. Dissecting a circuit for olfactory behaviour in Caenorhabditis elegans. Nature 450, 63–70 (2007).

38. Nagel, K. I. & Wilson, R. I. Mechanisms underlying population response dynamics in inhibitory interneurons of the Drosophila antennal lobe. J. Neurosci. 36, 4325–4338 (2016).

39. Nizampatnam, S., Zhang, L., Chandak, R., Li, J. & Raman, B. Invariant odor recognition with ON–OFF neural ensembles. Proc. Natl. Acad. Sci. 119, e2023340118 (2022).

40. Burton, S. D. Inhibitory circuits of the mammalian main olfactory bulb. J. Neurophysiol. 118, 2034–2051 (2017).

41. Chen, Z. & Padmanabhan, K. Top-down feedback enables flexible coding strategies in the olfactory cortex. Cell Rep. 38, 110545–110545 (2022).

42. Rothermel, M. & Wachowiak, M. Functional imaging of cortical feedback projections to the olfactory bulb. Front. Neural Circuits 8, 73 (2014).

43. Böhm, E., Brunert, D. & Rothermel, M. Input dependent modulation of olfactory bulb activity by HDB GABAergic projections. Sci. Rep. 10, 1–15 (2020).

44. Rothermel, M., Carey, R. M., Puche, A., Shipley, M. T. & Wachowiak, M. Cholinergic inputs from Basal forebrain add an excitatory bias to odor coding in the olfactory bulb. J. Neurosci. 34, 4654–4664 (2014).

45. Chandak, R. & Raman, B. Neural manifolds for odor-driven innate and acquired appetitive preferences. bioRxiv 2021.08.05.455310 (2021) doi:10.1101/2021.08.05.455310.

